# Host Switching Mutations in H5N1 Influenza Hemagglutinin Suppress Site-specific Activation Dynamics

**DOI:** 10.1101/2025.10.06.680362

**Authors:** Sally M. Kephart, Kiran F. Awatramani, Mason I. Saunders, Jacob T. Croft, Kelly. K. Lee

## Abstract

Increase in the occurrence of human H5N1 spillover infections resulting from dissemination of highly pathogenic avian influenza (HPAI) virus into bird and mammal populations raises concerns about HPAI adapting to become human transmissible. Studies identified hemagglutinin (HA) acid stability and receptor preference as essential traits that shape host tropism. Mutations that increase HA stability and affinity for α-2,6-linked sialic acids have been shown to confer airborne transmissibility in a ferret model, however mechanisms of activation of H5 subtype HA have not been probed and the effect of adaptive mutations on HA function has been largely inferred from static structures. Here, we use hydrogen/deuterium-exchange mass spectrometry to dissect activation dynamics for two ancestral HPAI H5 HA, their matched HA with adaptive mutations, and a contemporary H5 HA. By measuring dynamics, we identify variation in active site flexibility among the HA and demonstrate that adaptive mutations result in suppression of fusion peptide dynamics and stabilization of a key subunit interface involved in activation. The contemporary H5 isolated from a recent human spillover case exhibits a relatively protected fusion peptide and moderately depressed pH of activation compared to the HAs examined in this study. Our studies of activation dynamics in the H5 HAs in conjunction with prior analysis of H1 and H3 HA reveal subtype-specific patterns that correlate with adaptive mutation sites and indicate underlying physical constraints on influenza HA adaptation.

## Introduction

Highly pathogenic avian influenza (HPAI) viruses pose a persistent threat to ecosystems, economies, and human health. Currently, the impacts of HPAI are being felt across the globe due to widespread outbreaks of H5N1 virus. Since the virus emerged in late 2021 in North America, an unprecedented number of wild birds and poultry have died from infection and culling^1^. Frequent spillover of the virus into mammals has also meant devastation for other populations, including minks and seals^2^. For the first time, H5N1 also has been detected in dairy cows and in high titers in milk^3^. Notably, bovine mammary tissues have been shown to present both α-2,3 and α-2,6-linked sialic acid receptors, which support avian-origin virus replication while potentially facilitating adaptation to use of the α-2,6 receptor commonly associated with infection of humans^4,5^. The ability of the virus to spread between mammals raises concern that the barrier to human infection and transmission has been lowered. Indeed, a growing number of people have become infected with H5N1 after exposure to sick poultry and cows, but so far, no cases of sustained human-to-human transmission have been observed^2^. Studies of other ancestral HPAI A/H5N1 viruses suggest that further adaptation to humans would require changes to the trimeric hemagglutinin glycoprotein (HA) that mediates receptor binding and membrane fusion, as well as proteins involved in replication (PB2) and genomic segment packaging (NP) (Figure 1A)^6–12^. The experimental adaptation of reassortant H5N1 viruses has shown that both preferential binding to the human receptor and increased acid stability are required to confer airborne transmissibility in a ferret model (Figure 1B)^6,13^.

**Figure 1:**
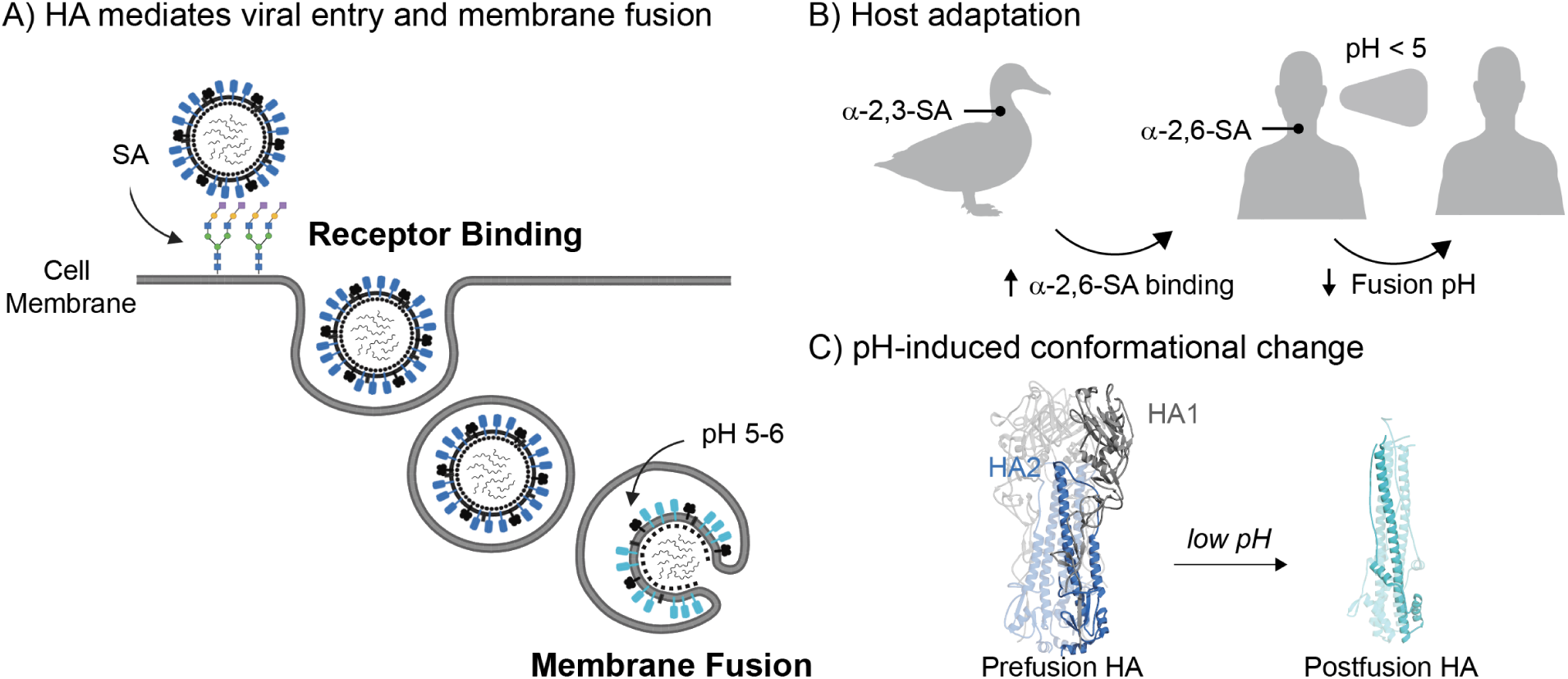
HA executes essential functions during the viral lifecycle and is a determinant of host tropism. **A)** HA (blue) binds to SA receptors on the surface of host cells, initiating viral entry. Acidification of the endosome triggers a cascade of structural changes in HA that drive fusion of the viral and host cell membranes. **B)** Adaptation to a human host requires changes to HA receptor specificity and acid stability. **C)** Low pH induces large-scale structural rearrangement in HA2 (blue) and dissociation of the HA1 subunits (grey) (PDB 6Y5G, 1QU1). Figure made with BioRender.

Residue substitutions associated with HA adaptation for airborne transmission have been identified for two strains of HPAI H5N1 virus, A/Vietnam/1203/2004 (Viet04) and A/Indonesia/5/2005 (Indo05)^6,7,13^. Comparing crystal structures of wildtype (WT) and mutant HA has revealed differences in the receptor binding domain (RBD) in the HA1 subunit including in positions directly lining the receptor binding site (RBS) that drive sialic acid (SA) preference, while mutations that are associated with increased acid stability in Viet04 and Indo05 have been identified at somewhat disparate sites along the HA1/HA2 interface (Figure 1C)^14–16^. The impact of acid-stabilizing mutations on mechanisms of HA activation and function have thus far remained unclear and are challenging to assess from available structural information^17–21^. Looking beyond static structures and probing activation with techniques that measure protein dynamics has revealed that the pathway from the prefusion to postfusion state involves a series of reversible transitions between highly dynamic intermediate conformations^22–30^. Understanding how acid-stabilizing mutations influence the activation process and fusion pathway is key to elucidating the physical constraints that shape how HA changes during adaptation to new hosts and routes of transmission.

Previously our lab has used hydrogen/deuterium exchange mass spectrometry (HDX-MS) to probe activation dynamics in HA from H1 and H3 influenza virus subtypes^22,25,26^. The studies identified trigger hotspots and specific regions of the protein that respond to changes in pH. Here, we apply a similar approach to study the effect of mutations that confer airborne transmissibility to HPAI H5N1 viruses on HA dynamics during acidification. Using the mutations identified for Viet04 and Indo05, we compare dynamics for WT and mutant HA across a range of pH values approaching the threshold for fusion. Our data shows that the adaptive mutations can alter RBS flexibility as well as suppressing dynamics in regions of the protein that respond to pH, however the effect on activation pathways is multi-faceted and there appear to be more than one mechanism for stabilizing HA while still allowing it to mediate pH activation and fusion. Namely, conformational change can be restrained by restricting fusion peptide release (Viet04) or by preventing head domain opening (Indo05). Additionally, we examine activation dynamics for HA derived from a contemporary H5N1 virus (A/Colorado/18/2022) providing insight into conservation and variation in H5 HA activation.

Elucidating the structural profile of activation dynamics in HA helps rationalize why adaptive mutations that have been documented emerge at specific interfaces and motifs in HA. Beyond H5N1, data aggregated from studies of activation dynamics in HA from different influenza virus subtypes (H1, H3, and now H5) reveal subtype-specific structural dynamic responses that coincide with the distribution of documented adaptive amino acid substitutions, thus providing a mechanistic rationale for the residue changes in HA that are observed to shift influenza virus host tropism and acid stability. This work thus provides a deeper understanding of physical constraints on influenza adaptation as we monitor the on-going evolution of HPAI H5N1 and influenza viruses.

## Results

### Mutations in Viet04 HA alter dynamics in the RBD and fusion domain

We first examined effects of the adaptive mutations identified by Imai *et al.* using soluble H5 constructs based on HA from A/Vietnam/CL26/2004 HPAI virus^6^. A combination of four mutations in the RBD and fusion domain was shown to confer airborne transmissibility in ferrets (N158D, N224K, Q226L, T318I; H3 numbering) (Figure 2A). In order to determine how the mutations impact prefusion HA dynamics, initial HDX-MS experiments were performed at pH 7.4. Our analysis highlights 79 homologous peptides that span 70% of the sequence. The same peptides could be identified under low pH conditions and in the other HA constructs examined in this study (Figure S1-4). Peptides consistently displayed unimodal HDX profiles, suggesting that the protein was conformationally uniform and there were no subpopulations of unprocessed or pre-triggered HA.

**Figure 2:**
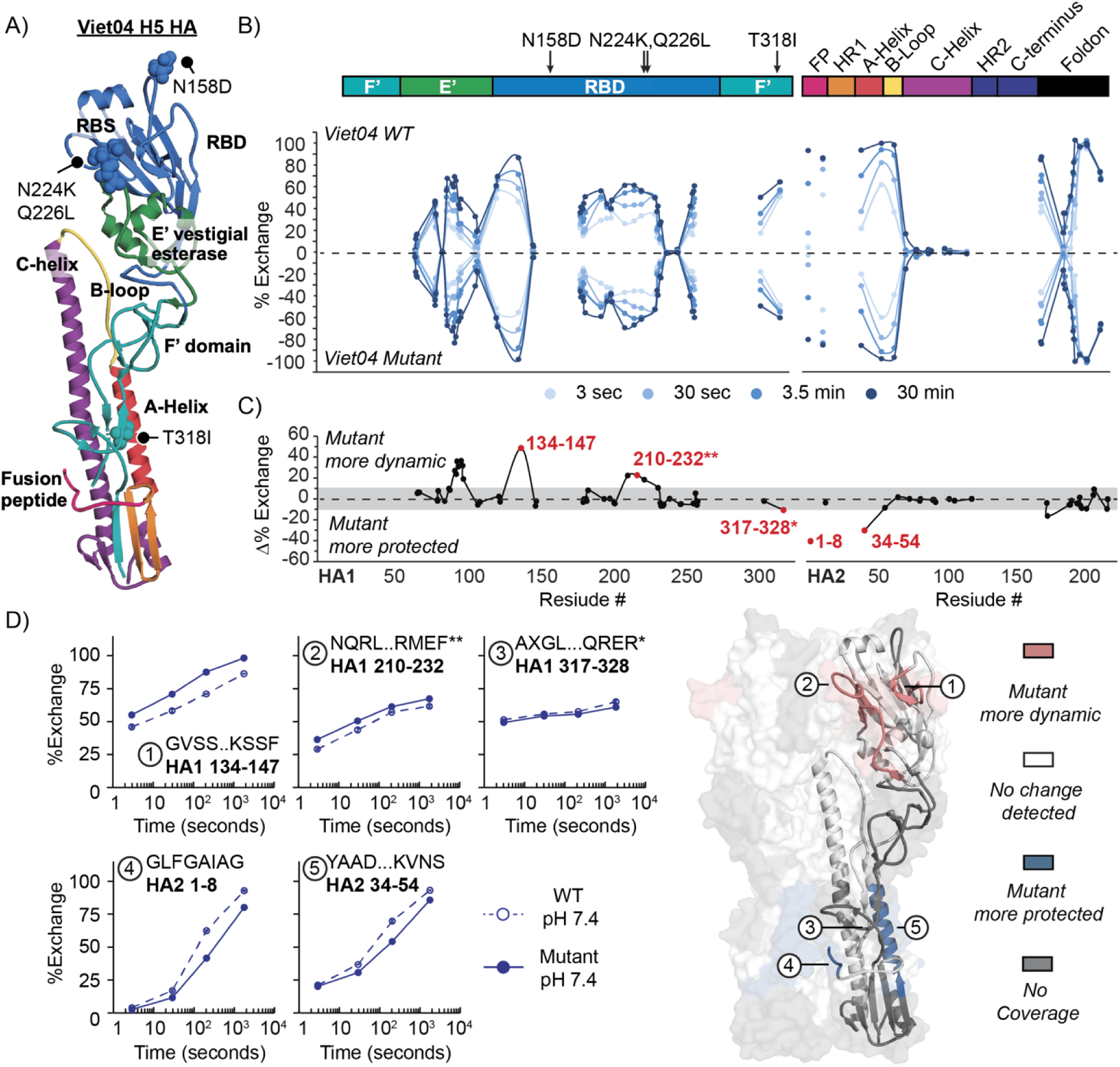
Mutations in Viet04 HA affect dynamics in RDB and fusion domain at neutral pH. **A)** Adaptive mutations (N158D, N224K, Q226L, T318I) mapped onto one protomer of Viet04 HA (PBD 4BGW) with domains colored: F’ domain (teal), vestigial esterase domain (E’; green), receptor binding domain (RBD; blue), fusion peptide (FP; pink), hairpin 1 (HR1; orange), A-helix (red), B-loop (yellow), C-helix (purple), hairpin 2 (HR2) and C-terminus (violet). **B)** Butterfly plots comparing exchange of WT (top) and mutant (bottom) at individual timepoints. The median residue position is plotted and aligned with the colored sequence map above. **C)** Residual plot showing differences between the WT and mutant proteins at neutral pH. Differences were determined using the total exchange for each peptide (% exchange summed across all timepoints). **D)** Differences greater than 10% are considered significant and are mapped onto HA structure (PDB 4BGW) with uptake plots shown for select peptides. Asterisks denote peptides containing mutations

While domain-level trends in H/D-exchange are similar for Viet04 WT and mutant (Figure 2B-C), differences in deuterium uptake between WT and mutant are observed in the RBS, specifically involving the 130- and 220-loops, which form two facets of the SA binding pocket and contain the sites of the N224K and Q226L mutations (Figure 2C). Peptides spanning these regions (Figure 2D; HA1 134-147, HA1 210-232) indicate that Viet04 mutant RBS in fact gains in dynamics relative to WT, while the third key motif involved in receptor binding, the 190-loop, exhibited nearly identical dynamic ordering in WT and mutant Viet04 HA. Increased dynamics in the receptor binding site is consistent with structural data that revealed that the receptor binding pocket is 1.5Å wider in Viet04 mutant^19,20^. The larger receptor binding pocket is associated with decreased affinity for the avian receptor and slightly increased affinity for the human receptor compared to WT, albeit the affinity for both receptors is relatively low^18–20^. Increased dynamics in the RBS may reflect changes in flexibility that enable greater receptor promiscuity. Differences in this region may also be related to the removal of a glycosylation sequon at position 158 in the mutant. Unfortunately, we lacked coverage of HA1 residues 155-176 and were not able to directly monitor changes near the site of N158D mutation.

Dynamic differences between Viet04 WT and mutant are also observed in the HA2 fusion subunit near the site of the acid-stabilizing mutation (T318I) (Figure 2D). Both the fusion peptide (HA2 1-8) and A-helix (HA2 34-54) display decreased dynamics in Viet04 mutant, which likely result from increased hydrophobic interactions with isoleucine at position 318 that stabilize the region^19,20^. Surprisingly, a peptide that contains the T318I site (HA1 317-328) shows minimal change in its dynamic profile, suggesting that stabilization in the fusion peptide and A-helix is due to the Ile sidechain and do not involve reorganization of the 317-328 peptide backbone.

### Mutations in Viet04 HA stabilize activation dynamics in HA2

To probe the effects of the mutations on HA activation, HDX reactions were also performed at acidic pH conditions approaching the threshold for fusion. The relevant pH for our experiments were determined using blue native PAGE (BN-PAGE) to visualize the formation of rosettes, which occurs at low pH in the absence of a membrane due to release of the hydrophobic fusion peptide^31^. For Viet04 WT, rosette formation (observed as HA retained in the wells) was first observed at pH 5.9, with high molecular weight oligomer formation increasing at pH 5.7 (Figure 3A). Equivalent trends were seen at pH 5.7 and 5.5 for Viet04 mutant, which is consistent with the mutant being more acid stable than WT (Figure 3A). HDX reactions were performed at the lowest pH conditions where HA remained well-behaved and soluble, pH 6.1 (WT) and 5.9 (mutant) (Figure 3B). In order to compare H/D-exchange at neutral and low pH, the deuteration time was scaled according to Li *et. al.*^32^. Internal exchange reporters included in the exchange reactions showed agreement across all conditions and confirm that the deuteration times used in the experiment were accurately matched (Figure S5)^33^.

**Figure 3:**
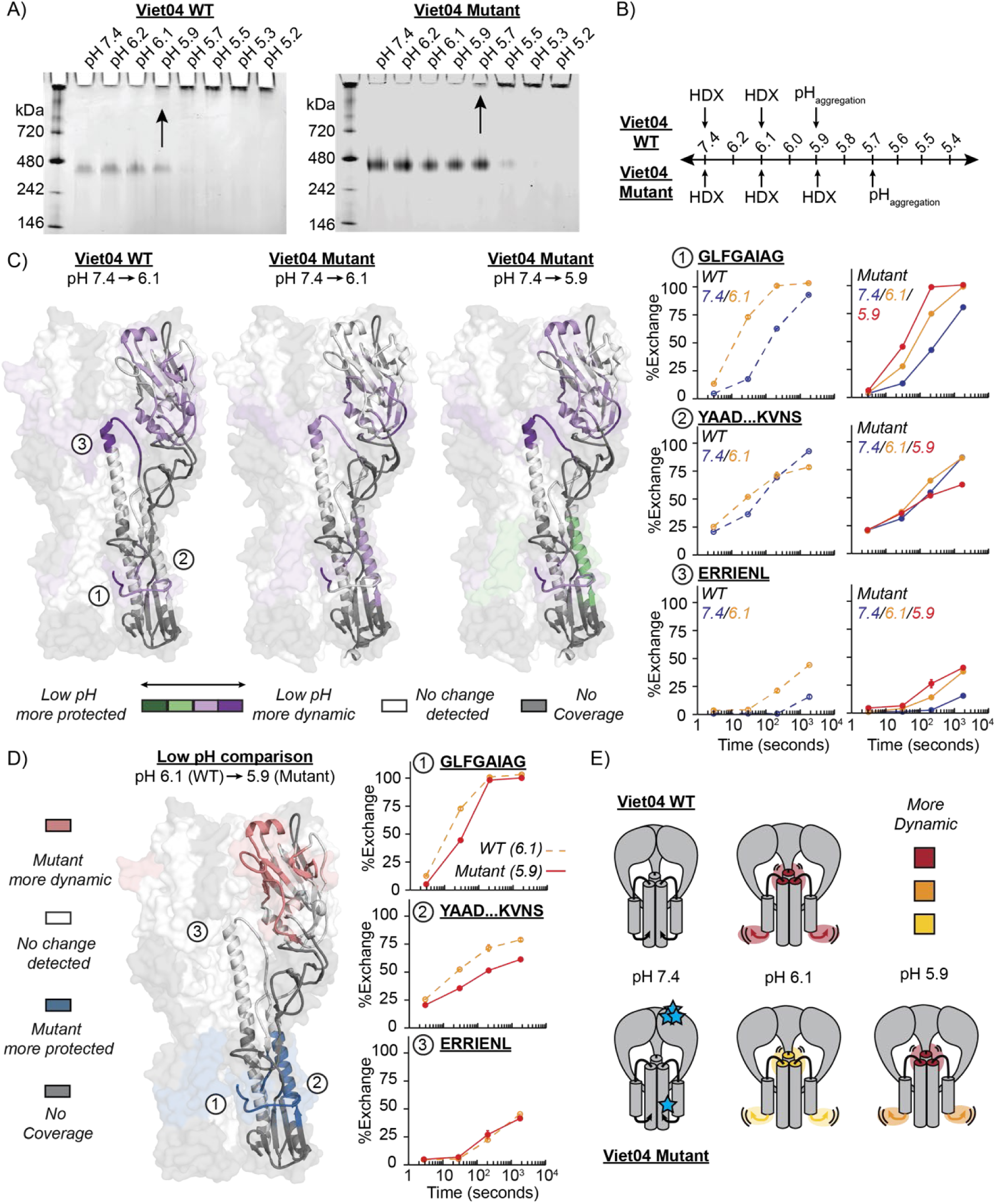
Mutations in Viet04 HA stabilize activation dynamics in HA2. **A)** BN-PAGE monitoring the formation of high molecular weight oligomers after 1 hour incubation at various pH. Arrows indicate the highest pH where aggregation was observed. **B)** Experiment overview. **C)** Differences in HA dynamics upon acidification are mapped onto the structures of Viet04 WT (PDB 4BGW) and mutant (PDB 4BH2). Differences were determined using the total exchange for each peptide (% exchange summed across all timepoints). Peptides that become more dynamic at low pH are colored purple and peptides that become more protected are colored green. The degree of shading reflects moderate (15-35%) and large (>35%) differences in total exchange. Differences totaling less than 15% are colored white, signifying no change. Peptides that could not be monitored due to low signal intensity or overlapping ions are colored grey. Uptake plots for three peptides are shown to right for WT (dashed lines) and mutant (solid lines). **D)** Differences in HA dynamics between Viet04 WT and mutant at the lowest pH where exchange was measured, pH 6.1 and pH 5.9 respectively, are mapped onto the structure of Viet04 WT (PDB 4BGW). Peptides that become more dynamic in the mutant are colored pink and peptides that become more protected in the mutant are colored blue. The same threshold for differences was applied as in part (C). Uptake plots for three peptides are shown to the right. **E)** Cartoon representation of activation dynamics for Viet04 WT (top) and mutant (bottom) showing degrees of fusion peptide release and head domain opening based on dynamic profile. Blue stars estimate the site of mutations.

Upon acidification, Viet04 WT displays increased dynamics throughout the trimer (Figure 3C, Table S1), most notably in the fusion peptide (Figure 3C, peptide 1), A-helix (Figure 3C, peptide 2), B-Loop, and top of the C-Helix at the HA1/HA2 interface (Figure 3C, peptide 3). Increased H/D-exchange in these regions indicates that the fusion peptide and HA1/HA2 interface are more exposed at pH 6.1 than pH 7.4. Exchanges at pH 6.1 were also performed on Viet04 mutant leading to increased dynamics throughout the trimer upon acidification including at the fusion peptide (Figure 3C, peptide 1) and top of the C-Helix (Figure 3C, peptide 3). However, comparing the activation profiles at pH 6.1 for Viet04 WT and mutant reveals that dynamic changes throughout the mutant HA are dampened relative to WT (Figure 3C, Table S1).

Further lowering the pH to 5.9 with the mutant increases H/D-exchange in many peptides compared to pH 6.1 resulting in an activation profile that nearly matches WT (Figure 3C, Table S1). Directly comparing deuterium uptake at the lowest pH condition reveals that dynamics are decreased in Viet04 mutant relative to WT near the site of the acid-stabilizing T318I mutation in the F’ domain, including in the fusion peptide and HR1/A-Helix (Figure 3C,D, peptides 1 and 2) (Table S3.1). Protection under these conditions indicates that activation dynamics related to fusion peptide release are restricted in the mutant even at the threshold of aggregation. Despite stabilization in the fusion subunit, many regions of the protein exchange similarly to WT at low pH, suggesting that other pH-responsive domains, such as the HA1/HA2 interface (Figure 3D, peptide 3), can be activated independently, thus HA in some regards behaves modularly rather than as a unitary assembly that exhibits long-range allosteric coupling (Figure 3E).

### Mutations in Indo05 HA affect dynamics at the HA1/HA2 interface at neutral pH

A minimal set of substitutions required to confer airborne transmissibility in ferrets has also been reported for HA from A/Indonesia/5/2005 HPAI virus (Figure 4A)^7,13^. Viet04 and Indo05 H5 HA share 96% sequence identity and the pattern of adaptive mutations show some parallels and differences for the two strains. Like Viet04, three mutations (T160A, Q226L, and G228S) are localized to the RBD in Indo05 and affect receptor preference. An acid-stabilizing mutation was also identified in Indo05, albeit in a different region than emerged in Viet04 HA. H110Y is located in the vestigial esterase (E’) domain at the HA1/HA2 interface in Indo05 HA. In order to determine whether different sets of mutations have distinct effects on HA dynamics, we repeated our HDX-MS experiments with soluble H5 constructs based on HA from A/Indonesia/5/2005 HPAI virus and the airborne transmission-adaptive mutations identified by Herfst *et al.* (Figure S3)^7^.

**Figure 4:**
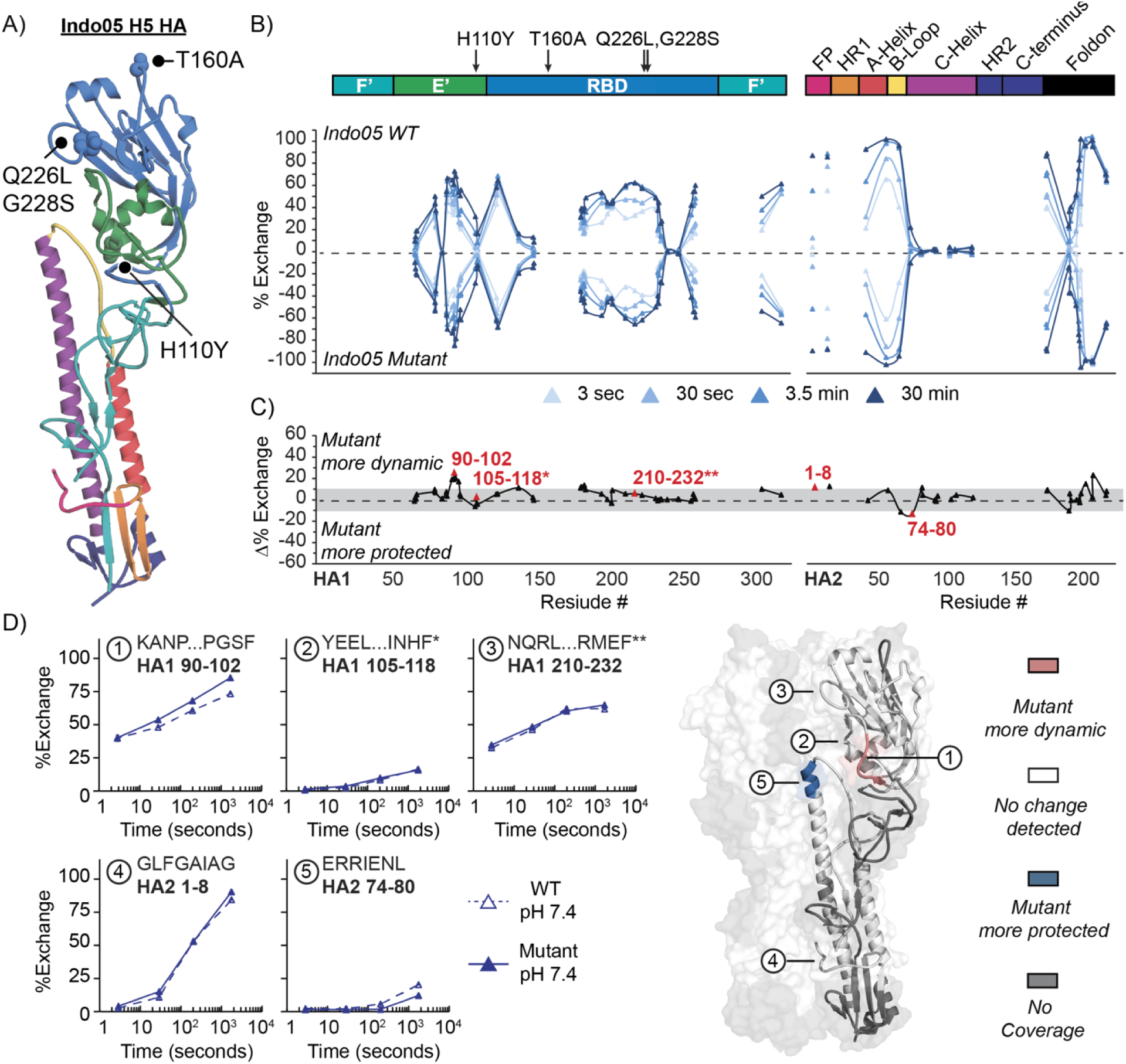
Mutations in Indo05 HA have a localized effect on dynamics at HA1/HA2 interface at neutral pH. **A)** Adaptive mutations (H110Y, T160A, Q226L, G228S) mapped onto one protomer of Indo05 HA (PBD 4K62) with domains colored similar to Figure 2. **B)** Butterfly plots comparing exchange of WT (top) and mutant (bottom) at individual timepoints. The median residue position is plotted and aligned with the colored sequence map above. **C)** Residual plot showing differences between the WT and mutant proteins at neutral pH. Differences were determined using the total exchange for each peptide (% exchange summed across all timepoints). **D)** Differences greater than 10% are considered significant and are mapped onto HA structure (PDB 4K62) with uptake plots shown for select peptides. Asterisks denote peptides containing mutations.

We initially performed HDX-MS experiments at pH 7.4 to determine how the mutations in Indo05 impact prefusion HA dynamics. Again, we observe that domain-level trends are generally similar for Indo05 WT and mutant (Figure 4B-C). In notable contrast to Viet04, the RBD of both WT and mutant Indo05 HA exhibited a considerably more ordered 130-loop than observed for Viet04 HA (Figure 4C, also see Figure 6C), while the 190- and 220-loops were similar in dynamic profile to Viet04 HA. Crystal structures show that, similar to Viet04, the receptor binding pocket in Indo05 mutant is 1Å larger than WT^21^. Surprisingly, in contrast to the reorganized Viet04 RBS, the adaptive residue changes that affect the 220-loop and glycosylation sequon at position 158 in Indo05 HA did not lead to increased dynamics in the mutant HA relative to WT in the receptor binding site or RBD more broadly (Figure 4C-D, e.g. peptide 3, HA1 210-232). HA1 peptide 90-102, which abuts residue 228, however exhibited elevated dynamics in response to the Q228S substitution, indicating a highly localized response to the adaptive mutations. Although both Viet04 and Indo05 mutant display increased binding to α-2,6 sialosides relative to wildtype, it has been shown that Viet04 mutant exhibits specificity for linear α-2,6 sialosides compared to Indo05 mutant, which binds to sialic acid on branched glycans^34^. This result supports our observation that the RBDs of Viet04 and Indo05 are structurally and dynamically distinct and are impacted in different ways by similar mutations.

In contrast to the relatively muted effects seen in the RBD, a notable decrease in H/D-exchange resulting from the transmission-adaptive mutations in Indo05 was observed proximal to the site of the H110Y acid-stabilizing mutation affecting the apex of C-helix in the HA2 central helical bundle (HA2 74-80). This HA2 peptide contains residue N79 (HA2), which forms a hydrogen bond with the Y110 (HA1) sidechain from the adjacent protomer in the mutant^21^. By contrast a peptide (HA1 105-118) itself directly spanning the 110-helix shows no difference in H/D-exchange, supporting the inference that the stabilization is mediated through the Y110 sidechain contacts. In contrast to the stabilization of the fusion peptide seen with the T318I mutation in Viet04, H110Y in Indo05 did not appear to alter fusion peptide (HA2 1-8) dynamics (Figure 4C,D).

### Mutations in Indo05 HA stabilize activation dynamics at the HA1/HA2 interface

To determine how the mutations identified for Indo05 H5N1 affect HA activation dynamics, HDX-MS experiments were also performed at acidic pH conditions approaching the threshold of activation. Using BN-PAGE, we observed that Indo05 WT and mutant started to aggregate into rosettes at pH values 5.9 and 5.7, respectively (Figure 5A). Delayed aggregation indicates that Indo05 mutant is more acid stable than WT. The shift in acid stability based upon the onset of low pH-induced aggregation between WT and mutant appears to be similar for Viet04 and Indo05 based on this assay and the conditions sampled, however we observed a notable persistence of soluble trimers even under low pH conditions for both WT and mutant Indo05 HA, suggesting the Indo05 HA trimers may maintain an prefusion-like organization well-below the threshold pH where some HA are being activated.

**Figure 5:**
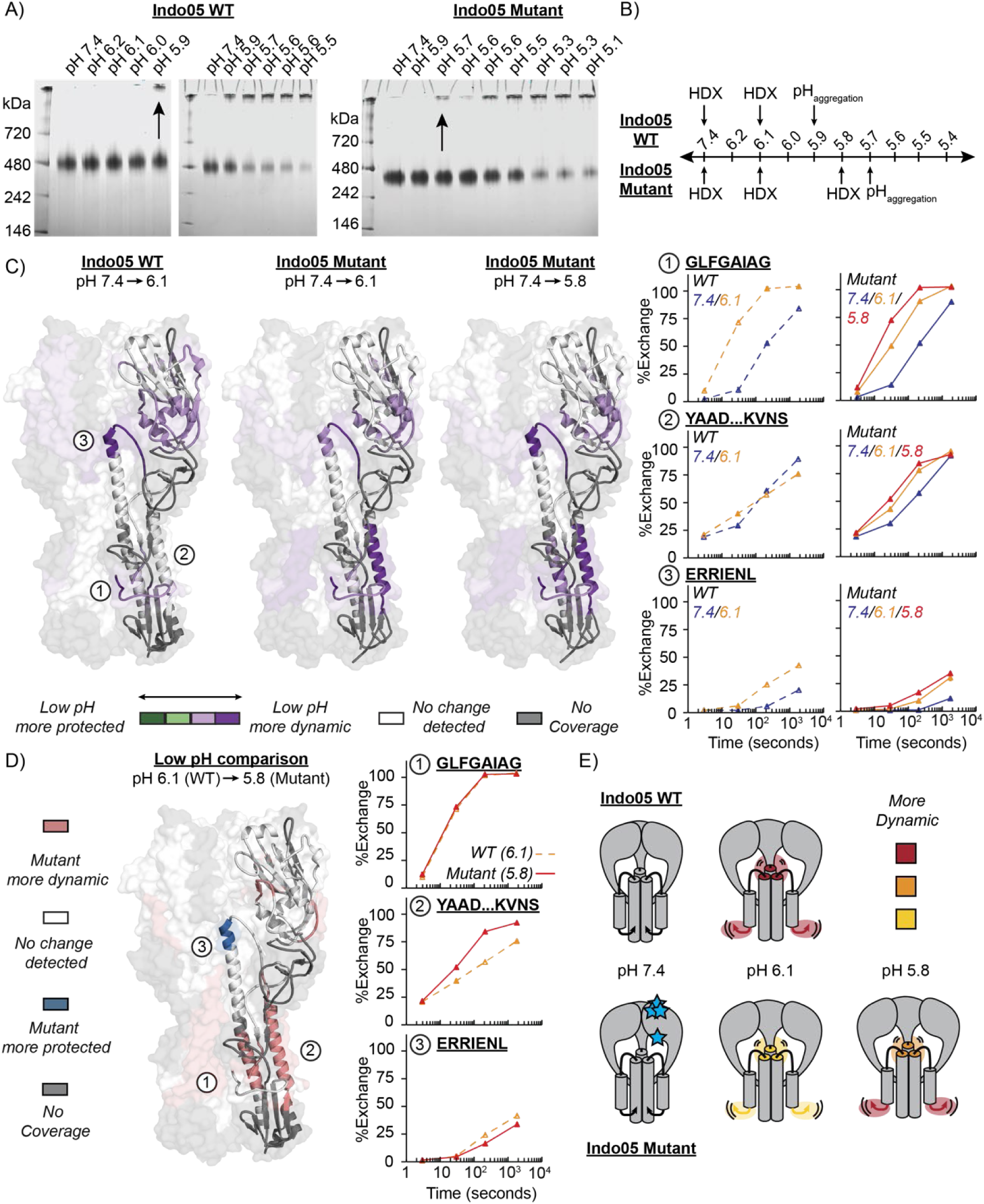
Mutations in Indo05 HA stabilize activation dynamics at the HA1/HA2 interface. **A)** BN-PAGE monitoring the formation of high molecular weight oligomers after 1 hour incubation at various pH. Arrows indicate the highest pH where aggregation was observed. **B)** Experiment overview. **C)** Differences in HA dynamics upon acidification are mapped onto the structures of Indo05 WT (PDB 4K62) and mutant (PDB 4K65). Differences were determined using the total exchange for each peptide (% exchange summed across all timepoints). Peptides that become more dynamic at low pH are colored purple and peptides that become more protected are colored green. The degree of shading reflects moderate (15-35%) and large (>35%) differences in total exchange. Differences totaling less than 15% are colored white, signifying no change. Peptides that could not be monitored due to low signal intensity or overlapping ions are colored grey. Uptake plots for three peptides are shown to right for WT (dashed lines) and mutant (solid lines). **D)** Differences in HA dynamics between Indo05 WT and mutant at the lowest pH where exchange was measured, pH 6.1 and pH 5.8 respectively, are mapped onto the structure of Indo05 WT (PDB 4K62). Peptides that become more dynamic in the mutant are colored pink and peptides that become more protected in the mutant are colored blue. The same threshold for differences was applied as in part (A). Uptake plots for three peptides are shown to the right. **E)** Cartoon representation of activation dynamics for Indo05 WT (top) and mutant (bottom) showing degrees of fusion peptide release and head domain opening based on dynamic profile. Blue stars estimate the site of mutations.

Exchanges were carried out at the lowest pH where HA remained well-behaved and did not show signs of aggregation, pH 6.1 (WT) and pH 5.8 (mutant) (Figure 5B). Similar to Viet04 WT, the activation profile of Indo05 WT captures changes in dynamics in the vestigial esterase and F’ domains of HA1 and the HA2 fusion subunit at pH 6.1, which contain clusters of pH-sensing residues (Figure 5C, Table S2)^14,15,35^. In HA2, increases in dynamics are observed in the fusion peptide (Figure 5C, peptide 1), B-Loop, and top of the C-Helix at the HA1/HA2 interface (Figure 5C, peptide 3). Similar regions are activated in Indo05 mutant at pH 6.1, although these regions do not become as dynamic as in WT at the same pH (Figure 5C, Table S2). The A-helix (Figure 5C peptide 2) by contrast exhibits greater dynamics in the mutant at pH 6.1.

Directly comparing deuterium uptake at the lowest pH conditions reveal differences between Indo05 WT and mutant at their respective thresholds of aggregation (6.1 for WT, 5.8 for mutant) (Figure 5D, Table S2). Significant protection is observed in the mutant at the top of the C-Helix, proximal to the site of acid-stabilizing mutation (H110Y) (Figure 5D, peptide 3). Decreased H/D-exchange at the HA1/HA2 interface could indicate that dynamics related to head domain opening are restricted in the mutant even at equivalent stages of activation. Despite stabilization in this region, the fusion subunit displays similar or increased dynamics compared to WT (Figure 5D, peptides 1-2). Notably the A-Helix (Figure 5D, peptide 2) in the Indo05 mutant remains more dynamic than in WT. This motif’s enhanced dynamics in Indo05 present a notable contrast to the stabilization seen in Viet04 mutant, in which the T318I mutation directly interacts with and stabilizes the A-helix (Figure 3D, peptide 2).

The contrasting modes of activation for acid-stabilized mutants, despite giving rise to similar shifts in pH of activation, demonstrate that effective stabilization can be take place in different ways resulting in different structural and dynamic effects on pathways of activation (Figure 5E, 3E).

### HA from a recent H5N1 HPAI avian-human spillover infection (A/Colorado/18/2022) exhibits a relatively protected fusion peptide and moderately depressed pH of activation

We sought to also characterize the dynamics and activation profile of H5 HA from a contemporary clade 2.3.4.4b H5N1 HPAI virus, which has been widespread in wild birds since 2021 and caused unprecedented outbreaks in poultry and mammals. Similar to Viet04 and Indo05, A/Colorado/18/2022 (Colo22) was isolated from a human who was exposed to sick poultry^36,37^. Colo22 H5 HA shares 92% sequence identity with Viet04 and Indo05 H5 HA and most sequence differences between the strains are localized to HA1 (Figure 6A).

**Figure 6:**
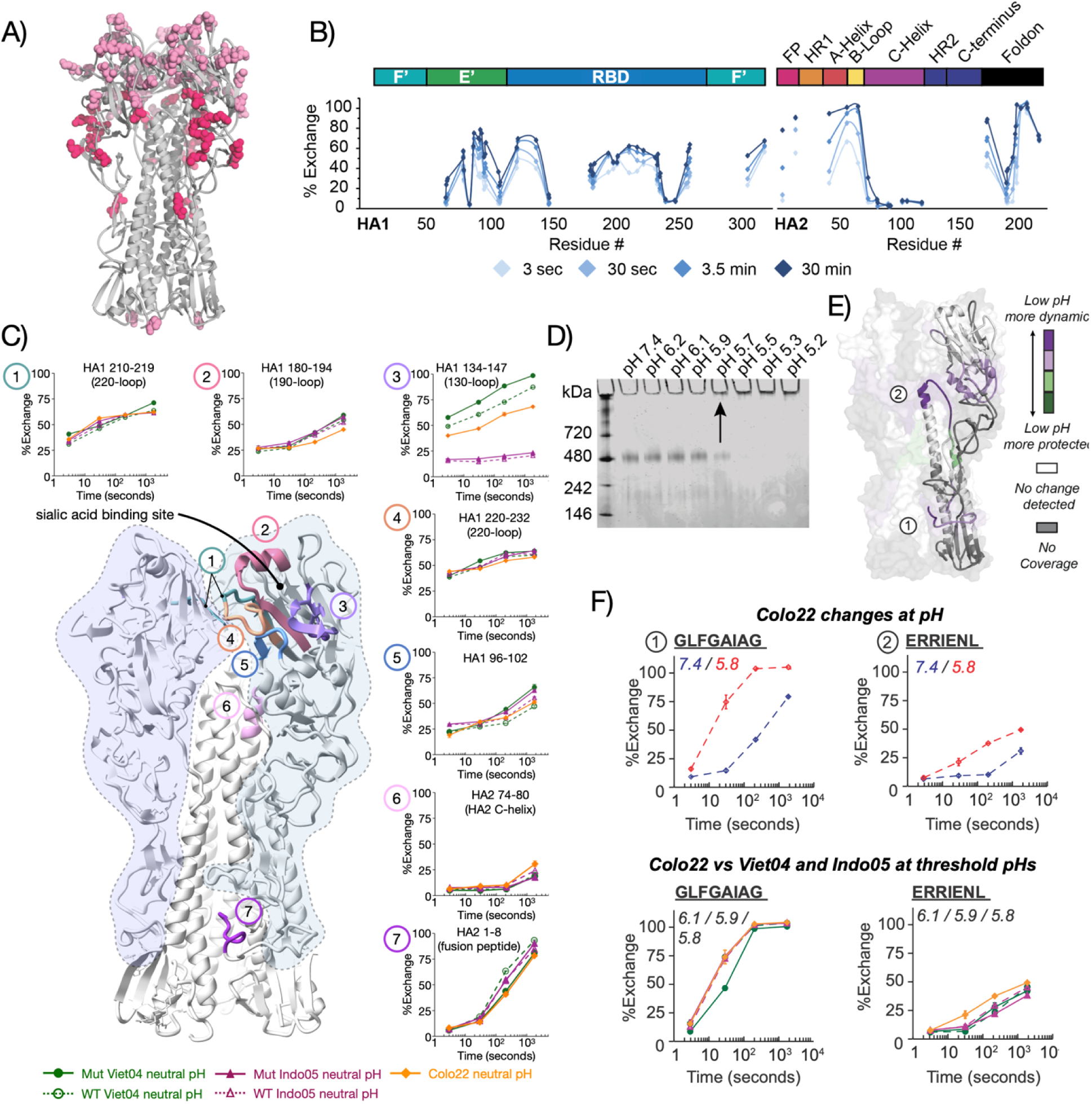
Colo22 displays similar activation profile to Viet04 and Indo05. **A)** Differences in Colo22 sequence (compared to Viet04 and Indo05 WT) are shown in pink. Substitutions that fall within pH-responsive regions are hot pink. B) Butterfly plot showing exchange at individual timepoints. The median residue position is plotted and aligned with the colored sequence map above. **C)** Structural dynamic variation in Colo22 receptor binding site and functional motifs versus Indo05 and Viet04 WT and mutant H5 HA. The sialic acid binding site, formed by multiple structural elements including the 130-, 150-, 190-, and 220-loops is a key site for adaptive mutation. HDX-MS reveals strain-specific variation as well as pH-dependent effects in a number of these motifs, with the 130-loop in particular exhibiting dramatic variation in dynamic flexibility between strains. In general, in response to low pH, the RBS motifs increase in dynamics. In order for all constructs to be compared, in-exchange was not corrected for due to Colo22 missing the 0 sec timepoint. **D)** BN-PAGE monitoring the formation of high molecular weight oligomers after 1 hour incubation at various pH. Arrows indicate the highest pH where aggregation was observed. Uptake plots comparing exchange at neutral pH across all constructs. **E)** Differences in HA dynamics upon acidification are mapped onto the structure of Viet04 WT (PDB 4BGW). Differences were determined using the total exchange for each peptide (% exchange summed across all timepoints). Peptides that become more dynamic at low pH are colored purple and peptides that become more protected are colored green. The degree of shading reflects moderate (15-35%) and large (>35%) differences in total exchange. Differences totaling less than 15% are colored white, signifying no change. Peptides that could not be monitored due to low signal intensity or overlapping ions are colored grey. Uptake plots for two peptides are shown to right. **F)** Uptake plots comparing exchange at low pH across all constructs. In the bottom panels, the uptake plots at each isolate’s respective pH threshold value are shown.

Comparing HDX-MS dynamic profiles at neutral pH for Colo22 (Figure 6B,C) with the other H5 HA we have examined, we observe that while dynamic profiles are generally similar across the set of HA, the RBS in Colo22 differs in ordering relative to Viet04 and Indo05 HAs, particularly involving the 130-loop (Figure 6C, peptide 3). In Colo22 the 130-loop exhibits dynamics intermediate between the highly ordered Indo05 and highly dynamic Viet04 HA (Figure 6C). RBS flexibility and conformational dynamics have been proposed to play a pivotal role in modulating receptor binding and promiscuity and, as we observed in Viet04, can change in response to acquisition of receptor switching mutations (Figure 2C,D). HDX-MS is thus an effective approach for quantifying the generally hard-to-measure traits of receptor binding site flexibility and loop ordering.

Colo22 HA also shows differences in other functionally important areas. For example, the fusion peptide in Colo22 is more protected at neutral pH than the other HA we examined, exchanging similarly to Viet04 mutant, which contains an acid-stabilizing mutation in the F’ domain that restricts fusion peptide release (Figure 6C). The peptide at the apex of the HA2 C-helix (HA2 74-80), which contacts the HA1 RBD 110-helix and connects to the HA2 B-loop was found to be moderately more dynamic in Colo22 relative to the other HA (Figure 6C, peptide 6).

We next examined whether the observed differences at neutral pH would impact how Colo22 responds to acidification. Using the BN-PAGE assay, we determined that Colo22 HA begins to show signs of activation around pH 5.7 (Figure 6C). Notably, syncytia formation has been reported under similar conditions in other recent studies of contemporary H5^38,39^. Importantly, by our BN-PAGE assay, Colo22 behaves more similarly to the Viet04 and Indo05 HAs that bear acid-stabilizing mutations. In addition, Colo22 exhibits a sharp response to low pH, as seen with Viet04 HA, rather than the more drawn-out pH response seen with Indo05 HAs. In order to further characterize the state of the HA trimers under the activation conditions, we performed negative stain electron microscopy, which demonstrated that the trimers exist either as individual prefusion-like HA trimers or as rosettes, which are associated by their hydrophobic fusion peptides (Figure S6)^40^.

HDX-MS was performed at pH 5.8 to capture dynamics at the threshold of Colo22 activation. The overall activation profile is broadly similar to the other H5 constructs with pH-linked changes in HA1 observed in the vestigial esterase E’ and F’ domains (Figure 6D). Additionally by HDX-MS, large increases in exchange at the fusion peptide and HA1/HA2 interface are observed (Figure 6D, peptides 1 and 2). Comparing deuterium uptake at low pH across all constructs reveals that the fusion peptide has adopted a dynamic configuration at its threshold of aggregation similar to the other HA except for Viet04 mutant, which has a highly protected fusion peptide (Figure 6F). This indicates that greater protection of the fusion peptide in Colo22 HA at neutral pH did not impact fusion peptide release during acidification.

In summary, we find that the Colo22 HA exhibits a similar profile of activation to the WT H5 HAs despite exhibiting a pH of activation ∼0.2 pH units lower than the Viet04 and Indo05 WT, which is in a similar range to the transmission adapted mutants.

## Discussion

Influenza HA acid stability and receptor preference are important molecular determinants of host tropism and viral phenotype. Many studies have identified mutations that shift the pH of fusion or change receptor preference; however, the underlying mechanisms that drive these effects have not always been clear from static structures or sequence information alone^41^. Acid-stabilizing mutations are especially challenging to study as they are thought to impact the highly dynamic, pH-dependent conformational change associated with fusion. Indeed, although they had been inferred to impact these dynamic processes, ours is the first study that demonstrates the structural and dynamic effects of the mutations on HA activation and function. Here, using HDX-MS, we compared HA from 3 “wild-type” H5N1 isolates and examined how the addition of mutations that confer airborne transmissibility impact HA activation. Without mutations, Viet04, Indo05, and Colo22 H5 HA generally display similar dynamics at neutral pH despite differences in their sequence, though with local variation observed at functionally critical sites including the HA1-HA1 interface, RBS, and fusion peptide regions.

### Variation in structural dynamics and flexibility are evident in the RBS among H5 HA

A number of studies have investigated RBS variation, many with a focus on sequence and conformational differences, often with receptors already bound^17,42–45^ The RBS is not only a functionally critical site, but is also a highly desirable target for neutralizing antibodies^46^. More recently, studies have put forward RBS structural flexibility and dynamics as a contributing trait that impacts sialic acid binding specificity and promiscuity^47,48^, however experimental approaches for assessing RBS flexibility have been lacking. We demonstrate that the majority of the structural motifs forming the RBS in the naturally occurring H5 HA (WT Viet04 and Indo05, and Colo22) are relatively conserved in their behavior, however the 130-loop exhibits dramatic dynamic variation among the H5 HAs (Figure 6C, peptide 3). Furthermore, at least in Viet04 130-loop flexibility also changes in response to the mutations that alter transmissibility and receptor specificity. A lesser degree of variation is observed for the 220-loop, which forms part of the HA1-HA1 interface. Of note, the adaptive mutations, particularly in the Viet04 background, led to a substantial increase in dynamics in both the 130- and 220-loops as well as in the peptide segment spanning from 96-102, which interacts with both the 220-loop and the apex of the HA2 C-helix (Figure 2C). In Viet04 HA, the increase in RBD dynamics resulting from the adaptive mutations was compensated by the T318I stabilizing mutation in the HA stem, leading to an HA trimer that exhibits a lower pH of activation even in the presence of elevated head domain dynamics and RBS flexibility.

### Distinct transmission-adaptive mutations result in different mechanisms of HA stabilization, impacting their mechanisms of activation

Despite the acquisition of similar adaptive mutations in the RBD, transmission-adapted Indo05 HA exhibited only modest changes in dynamics in its neutral pH, prefusion state with minor alteration of RBS structural dynamics (Figure 2). We infer that the H110Y mutation, which bolsters interactions with the HA2 C-helical apex (Figure 4C, HA2 peptide 74-80) and has been reported to add an H-bond to the C-helix^21^, results in a suppression of RBD destabilization. Previous studies have identified the 110-helix as a key motif whose interactions with the C-helix and B-loop in HA2 can have a significant modulatory effect on HA acid stability^49^. The distinct response to low pH seen in Indo05, however, is not solely a result of the H110Y mutation as the WT HA shows a persistent soluble trimer fraction even well-below the threshold for activation. The inclusion of H110Y and the other adaptive mutations extends this trend over an even broader pH range for the Indo05 mutant, whereas Viet04 WT and mutant as well as Colo22 HAs exhibit an abrupt transition between trimer and aggregate. The Indo05 BN-PAGE response is consistent with greater inherent stabilization of the RBD domain interactions, which must dissociate to uncage the HA2 fusion subunit when it is fully triggered^26,27,30^. Some influenza strains for example have been reported to exhibit inactivation pH values (where inactivation reflects irreversible transitioning to the postfusion form of HA) substantially lower than the activation pH values (reflecting fusion peptide release and initial conformational changes)^39,50,51^. H1 HA has been shown to have a high propensity to inactivate compared to H2 and H3^51^ and unlike H3, exhibits evidence of head domain opening at activation pH by HDX^22^, suggesting that HA1 dissociation affects HA stability at low pH more than does fusion peptide release.

In a previous investigation of the mutant residue contributions in Indo05 HA, Linster et al., observed that H110Y and the RBS mutations shifted pH of activation in opposing directions, resulting in a rebalancing of the mutant trimer’s pH response^13^. The H110Y substitution in Indo05 HA was speculated to have a similar effect to T318I in Viet04 due to both involving contacts with the HA2 fusion subunit, however we find that while both locally stabilize the trimers, the resulting effect on activation mechanisms for the Viet04 and Indo05 mutant HA are markedly different. While Viet04 locks down the fusion peptide proximal regions, the fusion peptide in Indo05 does not exhibit the same degree of restraint but instead exhibits a bolstered HA head assembly that contrasts with the Viet04 mutant whose head domains gain in dynamics and appear destabilized. In the H5 constructs we find the fusion peptides are relatively dynamic at neutral pH compared to other HA, with some such as Colo22 exhibiting measurably greater protection though still more dynamic than we have seen in an H3 HA (Aichi/68)^23^. These findings are consistent with single-molecule FRET data probing Viet04 WT HA dynamics on virions that demonstrated that the fusion peptide reversibly samples multiple states where it is released and displaced from the trimer core at neutral as well as at acidic pH ^30^.

### Subtype-specific profiles of activation dynamics and sites of acid-stabilization

Taken together, our results show that acid-stabilizing mutations decrease dynamics in pH-responsive regions during HA activation and shift the pH where membrane-active conformations are first observed. We compared these regions with sites where stabilizing mutations have been reported for H5 HA including residue substitutions identified in a recent extensive study by Dadonaite et al. where pseudoviruses bearing deep mutational scanning libraries of HA based on WHO-recommended H5 candidate vaccine strain were selected for 2-6 sialic acid usage and separately for resistance to acid inactivation (A/South Carolina/USDA-000345-001/2021)^6,13,42,49,52^. The sites identified as helping to stabilize HA against low pH were mapped in Figure 7A are found to overlap with the regions we have identified as changing in dynamics in the H5 HA examined in the present study. Even more generally, when a similar comparison of acid-stabilizing mutations identified in H1^53–58^ and H3^14,59,60^ HA’s is compiled, the distribution of those sites was found to overlap with regions where dynamic changes have been observed by HDX-MS in H1 and H3 HA respectively. Each subtype appears to exhibit a distinct overall profile of acid activation (Figure 7A), and adaptive mutations that stabilize the HA from that subtype emerge in the regions of dynamic activation to suppress dynamics, rendering the mutant virus less susceptible to premature activation and inactivation (Figure 7B)^22^. Compared to HA from H1 and H3 subtypes studied by HDX-MS ^22,25,26^, H5 is distinct in that it appears to exhibit concerted changes to the head domain interfaces, concentrated in the E’ vestigial esterase domain, as well as fusion peptide proximal region during activation (Figure 7A). However, of these, it is more similar to H1 HA, which belongs to the same phylogenetic group 1, while H3 HA exhibits a more staged process in which the head domain association is bolstered while the fusion subunit gains in dynamics^22,25^.

**Figure 7:**
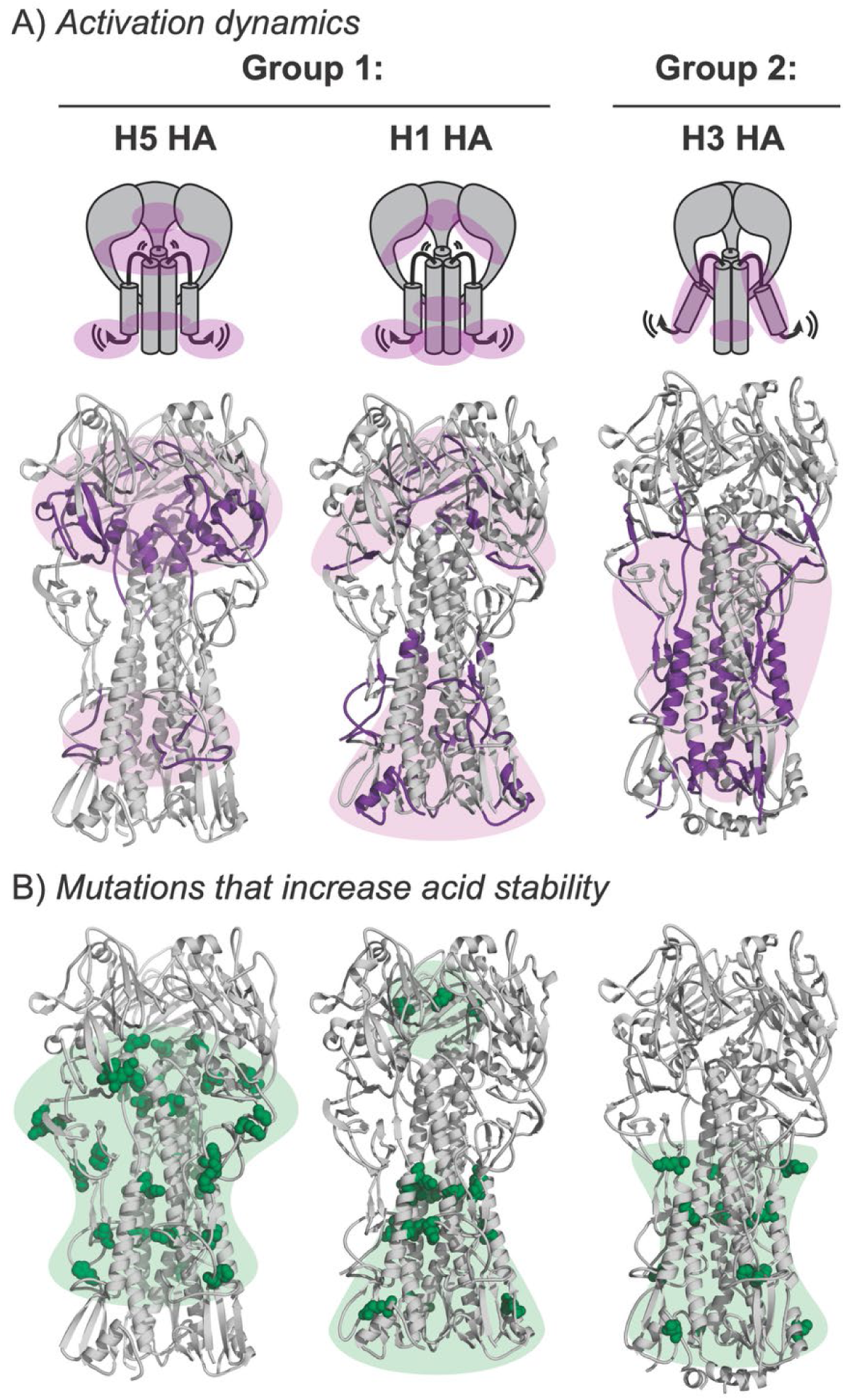
Mutations that modulate acid stability localize to pH-responsive regions in a subtype-specific way. **A)** Peptides that exhibit dynamic differences upon acidification for H5 (Viet04 WT, Indo05 WT, Colo22; PDB 4BGW), H1 (Puerto Rico/1934; PDB 1RU7), and H3 (Aichi/1968; PDB 6Y5G) are colored purple. **B)** Mutations that modulate HA acid stability are shown in green ^6,13,42,49,52,53–58,14,59,60^. For H5, dynamics and acid shifting mutations fall within the HA stalk and fusion peptide proximal region as well as in the F’ and vestigial esterase E’ domain including contact sites with the apex of the HA2 C-helix bundle (see structural motif annotations in Figure 2A). For H1 HA, mutations at the HA1-HA1 interface are seen, which also undergoes dynamic uncaging^22^ as well as in the HA stalk A-helix region and stalk base. By contrast, in H3 HA, the HA1-HA1 interactions remain stable and become more ordered approaching the threshold of activation^22,25^, but dynamics and acid stabilizing mutations cluster around the HA2 stalk regions, which respond first to low pH.

These studies of influenza HA activation dynamics provide mechanistic insight for why they adaptive changes emerge where they do in each subtype. We believe this is a powerful framework for understanding mechanisms of host adaptation because it reveals how HA changes structurally and functionally and suggests that differences in activation dynamics may underlie the variable effect of acid-stabilizing mutations across strains and subtypes^41,61^. Likewise, these new findings advance our understanding of viral fusion glycoprotein function and can help inform engineering of more stable HA-based immunogens for vaccines.

## Methods

### Cloning and mutagenesis

Sequences for the HA ectodomain (residues 11-329 of HA1 and 1-176 of HA2, H3 numbering) from A/Vietnam/CL26/2004, A/Indonesia/5/2005, and A/Colorado/18/2022 were codon optimized for mammalian expression and synthesized as gBlocks (Integrated DNA Technologies). Sequences included the native signaling peptide and polybasic cleavage site. The gBlocks were cloned into the CMV/R (VRC 8400) mammalian expression vector, which was kindly provided by Dr. Masaru Kanekiyo and contains a C-terminal foldon trimerization domain, Avi-tag, and hexa-histidine tag. When desired, mutations were added using the Q5 site-directed mutagenesis kit from New England Biolabs. Sequences were confirmed by whole-plasmid sequencing (Primordium).

### Mammalian expression and purification of HA

Recombinant expression of HA was carried out in Expi293F suspension cells cultured in Freestyle 293 Expression medium maintained at 37°C and 8% CO_2_. Cells were transiently transfected at a density between 2.5-3.0 million cells/mL using polyethylenimine (PEI) and Opti-MEM (Gibco) with HA and furin plasmids. For all constructs except Colo22, the ratio of HA to furin DNA was 3:1, which consistently generated cleaved HA1/HA2 based on SDS-PAGE. For Colo22, the ratio was decreased to 2.5:1, which yielded similar results based on SDS-PAGE. Cells were harvested 4-5 days post transfection when they were roughly 60% viable (Figure S7A). The supernatant was collected by centrifugation and filtered through a 0.45-μM PES filter (Nalgene). Tris-HCl pH 8 was added to 20 mM final concentration, along with 0.02% sodium azide (final concentration), and protease inhibitors (Pierce). The supernatant was transferred to a roller bottle with 1 mL Ni-NTA resin (Thermo Fisher Scientific) for overnight batch-binding at 4°C. The following day, the resin was collected in a gravity flow column at room temperature and washed with 30 column volumes of wash buffer (20 mM Tris-HCl [pH 8.0], 0.5 M NaCl, 10 mM imidazole, 0.02% sodium azide). The bound protein was eluted with 5 column volumes of elution buffer (20 mM Tris-HCl [pH 8.0], 0.5 M NaCl, 0.5 M imidazole, 0.02% sodium azide) five times. All fractions (flow-through, wash, and elution) were analyzed by SDS-PAGE. Elution fractions containing HA were combined, buffer exchanged into HEPES buffered saline (HBS; 150 mM NaCl, 10 mM HEPES [pH 7.4], 0.02% sodium azide), and concentrated using a 10K spin concentrator (Vivaspin, 20 mL, Sartorius).

For ion-exchange chromatography, the protein was dialyzed overnight into low salt buffer (20 mM Tris-HCl [pH 8.0], 1 mM EDTA) using 10K SnakeSkin dialysis tubing (Thermo Scientific) and then loaded onto a 1-mL HiTrapQ column (Cytiva). After 10 minutes of isocratic flow in 100 mM NaCl, a linear gradient up to 1 M NaCl was initiated. HA consistently eluted in 30-40 mM NaCl.

Prior to size-exclusion chromatography (SEC), the sample was treated with neuraminidase (NA) to prevent inter-trimer interactions that were thought to be driven by NeuAc glycans present on HA (Figure S8). For the NA treatment, 1 enzyme unit (EU) of α-3,6,8 neuraminidase (New England Biolabs) was added per 1 μg of HA and incubated at 37°C for one hour with rocking. Following treatment, the protein was concentrated to approximately 0.5 mL using a 10K spin concentrator (Amicon, 0.5 mL, Millipore) and then injected onto a Superdex 200 Increase 10/300 GL (Cytiva) column using an AKTA Pure (GE) system flowing HEPES buffered saline (HBS; 150 mM NaCl, 10 mM HEPES [pH 7.4], 0.02% sodium azide) at 0.5 mL/min. Trimeric HA eluted between 10.1 and 10.5 minutes for all constructs (Figure S7B).

### SDS-PAGE and BN-PAGE

SDS-PAGE and blue native PAGE (BN-PAGE) were performed to assess sample purity throughout the purification and prior to experiments (Figure S6C). For SDS-PAGE, samples were heated for 10 minutes at 90°C to ensure full denaturation. Reduced samples were prepared using 60 mM dithiothreitol (DTT). BN-PAGE was also used to assess pH-dependent aggregation of HA. For these experiments, purified HA in HEPES buffered saline (HBS; 150 mM NaCl, 10 mM HEPES [pH 7.4], 0.02% sodium azide) was diluted 1:1 in acidification buffer (75 mM NaCl, 10 mM Na_3_C_6_H_5_O_7,_ 10 mM HEPES, 0.02% sodium azide). Separate acidification buffers were prepared to sample a range of pH between 5 and 7.4. The pH values listed in the paper reflect the final pH of the diluted sample. After dilution, HA was incubated for 1 hour at room temperature prior to running the gel.

### Hydrogen/deuterium exchange

Samples were prepared using 5 μg of HA incubated in deuterated buffer (150 mM NaCl, 10 mM Na_3_C_6_H_5_O_7,_ 10 mM HEPES, 0.02% sodium azide, 85% D_2_O, pH* 7.4) for 3 seconds, 30 seconds, 3.5 minutes, and 30 minutes at room temperature. The reaction was stopped by adding an equal volume of ice-cold quench buffer (2 M urea, 200 mM tris(2-chorethyl) phosphate (TCEP), 0.2% formic acid) to a final pH of 2.5. The samples were immediately frozen with liquid nitrogen and stored at −80°C until analysis. All exchanges were performed in triplicate. Low pH exchanges were carried out as described above, but the deuteration time was adjusted to account for the effect of pH on the intrinsic exchange rate. Equivalent deuteration times for pH* 6.1 and 5.8 were determined using Equation 1, assuming a 10-fold decrease in exchange for every 1 unit decrease in pH* ^32^. Precise pH* measurements and labeling times are listed in Table S5. Internal exchange reporters were provided by Dr. Taylor Murphree and included in each reaction to ensure that conditions were consistent throughout the experiment (Figure S5)^33^.

Undeuterated controls were prepared similarly but using optima water in place of D_2_O. The zero timepoint was made by adding the protein directly to quenched deuteration buffer before freezing. Fully deuterated controls were prepared for each construct by collecting pepsin digestion eluate from two undeuterated samples. The digested peptides were concentrated under vacuum, resuspended in deuteration buffer, and heated to 65°C for one hour before they were subsequently quenched and frozen. To monitor protein stability during the longest exchange reaction, the equivalent of a 3 second pulse was performed alongside the adjusted 30 minute timepoint for the pH* 5.8 sample.

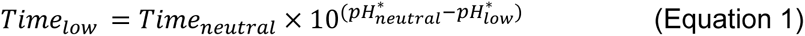

### Mass spectrometry and data analysis

Samples were thawed on ice and manually injected onto an online pepsin column (2.1 × 50 mm) kept at 15°C while flowing loading buffer (2% acetonitrile, 0.1% trifluoroacetic acid) at 400 μL/min^62^. Digested peptides were trapped on a CSH C18 trap cartridge (2.1 × 50 mm, Waters) for three minutes and resolved over an ACQUITY UPLC CSH C18 VanGuard column (130 Å, 1.7 µm, 1 × 100 mm, Waters) using a 15 minute linear gradient from 3% to 40% B (A: 98% water, 2% acetonitrile, 0.1% formic acid, 0.025% trifluoroacetic acid; B: 100% acetonitrile, 0.1% formic acid; flow rate: 40 μL/min). The column and plumbing were housed in a custom cold box kept at 0°C^62^. Data was analyzed on a Synapt G2-Si Q-TOF (Waters) mass spectrometer with the StepWave ion optics settings adjusted to minimize deuteration loss during ionization, and source and desolvation temperatures set to 70°C and 130°C, respectively^63^. A series of wash steps were implemented between injections to prevent carryover^64^.

Peptide assignments were made from MS^E^ data collected for undeuterated samples using Byonic (Version 3.8, Protein Metrics Inc.). Peptides were identified from DriftScope (Waters) using retention time and drift time coordinates. Deuterium uptake analysis was performed with HD-Examiner (Sierra Analytics) and HX-Express v2 for binomial fitting^65,66^. Percent exchange was determined using the fully deuterated control for each construct. When possible, in-exchange was corrected for using the 0 timepoint. Data for each timepoint was collected in duplicate. Pulse controls for pH 5.8 were included as a third replicate for the 3 sec timepoint. Experimental error was calculated using the standard deviation from duplicate measurements. PyMOL (Schrodinger, LLC) was used to generate figures.

### Negative stain electron microscopy

Purified Colo22 HA (0.02 mg/mL) in HEPES buffered saline (10 mM HEPES, 150 mM NaCl, 0.02% sodium azide) was diluted 1:1 with acidification buffer (10 mM HEPES, 10 mM citrate, 150 mM NaCl, 0.02% sodium azide) and allowed to incubate at 22 °C for 1 hour at each pH 7.4 or 5.6. At the hour mark, 3 uL of each sample was applied for 1 minute to a glow discharged carbon film 300 mesh negative stain grid (Quantifoil; Electron Microscopy Sciences). The sample was then blotted to dryness, washed with DI water, blotted again to dryness, and then a 2% solution of methylammonium tungstate (NanoW^TM^; Electron Microscopy Sciences) was applied for 1 minute before finally blotting to dryness. Micrographs were collected on an FEI Tecnai G2 Spirit 120 kV TEM using a Gatan Ultrascan 4000 4k x 4k CCD Camera System with a pixel size of 1.60 Å /px, a nominal dose of 35 – 40 e^−^/Å^2^, and defocus set to −2.00 µm using Leginon EM acquisition software^67^. Brightness and contrast of representative micrographs were auto-adjusted using Fiji and an 8 pixel median filter^68^.

## Data availability

According to community-based recommendations, HDX summary tables and uptake plots are included in the supplement (Tables S1-3, S5 and Figures S2-4)^69^. The mass spectrometry proteomics data have been deposited to the ProteomeXchange Consortium via the PRIDE^70^ partner repository with the dataset identifier PXD059353. Additional data supporting the findings of this paper are available from corresponding authors upon reasonable request.

## Supporting information

This article contains supporting information.

## Acknowledgements

We thank Professors Miklos Guttman, Abhinav Nath, Neil King, and Sharona Gordon for thoughtful discussions and Dale Whittington from the University of Washington School of Pharmacy Mass Spectrometer Center for help and support.

