## Supplemental figures and tables for "Host Switching Mutations in H5N1 Influenza Hemagglutinin Suppress Site-specific Activation Dynamics"

**HA1**

11 … 54a … 82a 88
**Viet2004** DQICIGYHANNSTEQVDTIMEKNVTVTHAQDILEKTHNGKLCDLDGVKPLILRDCSVAGWLLGNPMCDEFINVPEWSYIV
**Indo2005** DQICIGYHANNSTEQVDTIMEKNVTVTHAQDILEKTHNGKLCDLDGVKPLILRDCSVAGWLLGNPMCDEFINVPEWSYIV
**Colo2022** DQICIGYHANNSTEQVDTIMEKNVTVTHAQDILEKTHNGKLCDLNGVKPLILKDCSVAGWLLGNPMCDEFIRVPEWSYIV

89 92a … H110Y … 125a-b … 133a … N158D T160A 164
**Viet2004** EKANPVNDLCYPGDFNDYEELKHLLSRINHFEKIQIIPKSSWSSHEASLGVSSACPYQGKSSFFRNVVWLIKKNSTYPTI
**Indo2005** EKANPTNDLCYPGSFNDYEELKHLLSRINHFEKIQIIPKSSWSDHEASSGVSSACPYLGSPSFFRNVVWLIKKNSTYPTI
**Colo2022** ERANPANDLCYPGSLNDYEELKHMLSRINHFEKILIIPKSSWPNHETSLGVSAACPYQGAPSFFRNVVWLIKKNDAYPTI

165 … N224K Q226L G228S … 244
**Viet2004** KRSYNNTNQEDLLVMWGIHHPNDAAEQTKLYQNPTTYISVGTSTLNQRLVPRIATRSKVNGQSGRMEFFWTILKPNDAIN
**Indo2005** KKSYNNTNQEDLLVLWGIHHPNDAAEQTRLYQNPTTYISIGTSTLNQRLVPKIATRSKVNGQSGRMEFFWTILKPNDAIN
**Colo2022** KISYNNTNREDLLILWGIHHSNNAEEQTNLYKNPTTYISVGTSTLNQRLAPKIATRSQVNGQRGRMDFFWTILKPDDAIH

245 … 262a … T318I 323
**Viet2004** FESNGNFIAPEYAYKIVKKGDSTIMKSELEYGNCNTKCQTPMGAINSSMPFHNIHPLTIGECPKYVKSNRLVLATGLRNS
**Indo2005** FESNGNFIAPEYAYKIVKKGDSAIMKSELEYGNCNTKCQTPMGAINSSMPFHNIHPLTIGECPKYVKSNRLVLATGLRNS
**Colo2022** FESNGNFIAPEYAYKIVKKGDSTIMKSGVEYGHCNTKCQTPVGAINSSMPFHNIHPLTIGECPKYVKSNKLVLATGLRNS

324 328a-d 329
**Viet2004** PQRERRRKKR
**Indo2005** PQRESRRKKR
**Colo2022** PLREKRRK-R

**HA2**

1 … 80
**Viet2004** GLFGAIAGFIEGGWQGMVDGWYGYHHSNEQGSGYAADKESTQKAIDGVTNKVNSIIDKMNTQFEAVGREFNNLERRIENL
**Indo2005** GLFGAIAGFIEGGWQGMVDGWYGYHHSNEQGSGYAADKESTQKAIDGVTNKVNSIIDKMNTQFEAVGREFNNLERRIENL
**Colo2022** GLFGAIAGFIEGGWQGMVDGWYGYHHSNEQGSGYAADKESTQKAIDGVTNKVNSIIDKMNTQFEAVGREFNNLERRIENL

81 … 160
**Viet2004** NKKMEDGFLDVWTYNAELLVLMENERTLDFHDSNVKNLYDKVRLQLRDNAKELGNGCFEFYHKCDNECMESVRNGTYDYP
**Indo2005** NKKMEDGFLDVWTYNAELLVLMENERTLDFHDSNVKNLYDKVRLQLRDNAKELGNGCFEFYHKCDNECMESIRNGTYNYP
**Colo2022** NKKMEDGFLDVWTYNAELLVLMENERTLDFHDSNVKNLYDKVRLQLRDNAKELGNGCFEFYHKCDNECMESVRNGTYDYS

161 … 176 … 229
**Viet2004** QYSEEARLKREEISGV**GSGYIPEAPRDGQAYVRKDGEWVLLSTFLGSGLNDIFEAQKIEWHEGHHHHHH**
**Indo2005** QYSEEARLKREEISGV**GSGYIPEAPRDGQAYVRKDGEWVLLSTFLGSGLNDIFEAQKIEWHEGHHHHHH**
**Colo2022** QYSEEARLKREEISGV**GSGYIPEAPRDGQAYVRKDGEWVLLSTFLGSGLNDIFEAQKIEWHEGHHHHHH**

Number of Peptides: 84
Average Peptide Length: 11.2 residues
Coverage: 70%
Redundancy: 1.6

**Figure S1: HDX-MS sequence coverage**
Mature sequences for HA constructs are listed and numbered according to the H3 convention. Appended sequences including the foldon trimerization domain, Avi tag, and hexa-histidine tag

are bolded. Host-adaptation mutations in Viet04 and Indo05 are shown in red. Peptides where HDX was monitored are underlined.

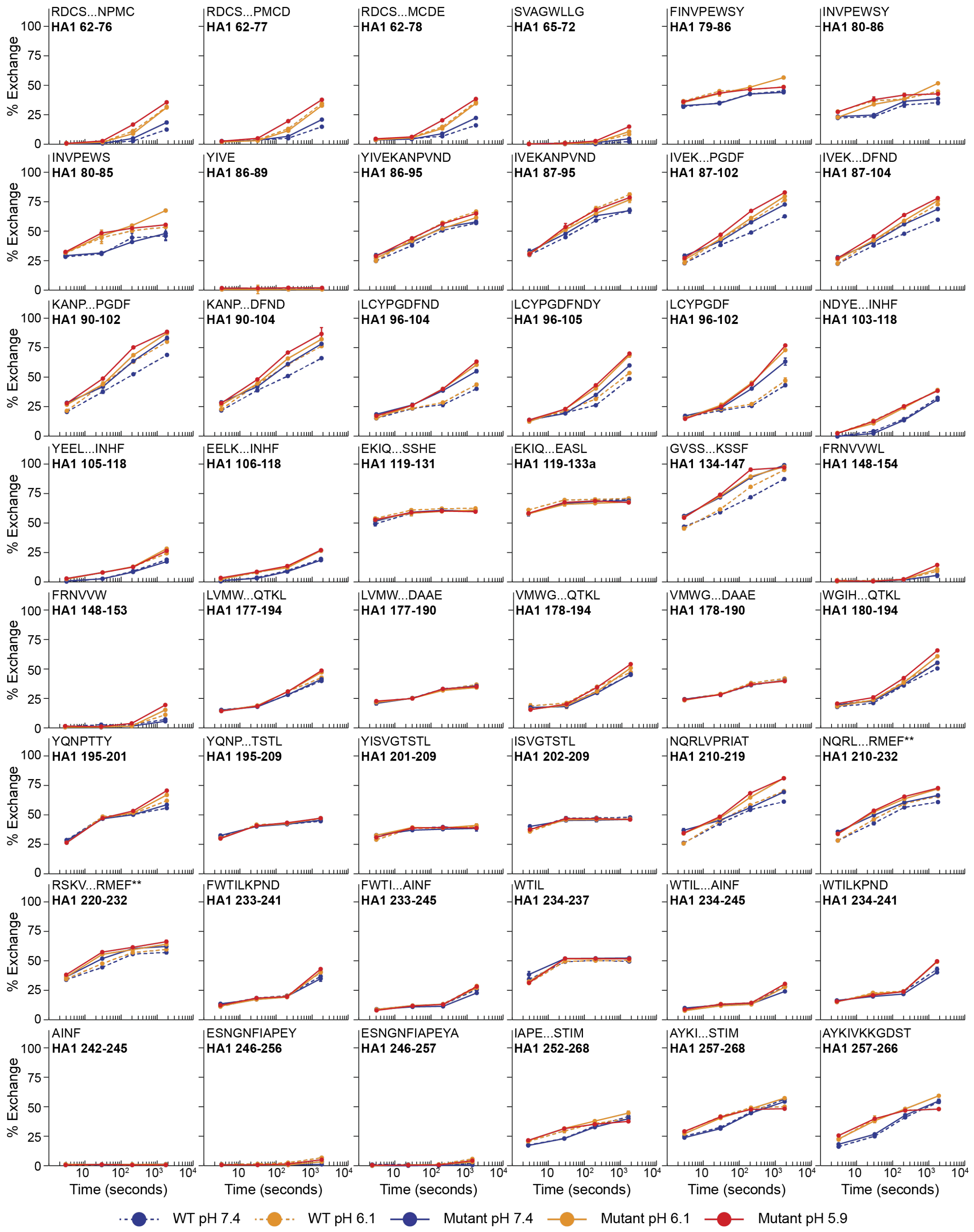
RDCS...NPMC
HA1 62-76
RDCS...PMCD
HA1 62-77
RDCS...MCDE
HA1 62-78
SVAGWLLG
HA1 65-72
FINVPEWSY
HA1 79-86
INVPEWSY
HA1 80-86
INVPEWS
HA1 80-85
YIVE
HA1 86-89
YIVEKANPVND
HA1 86-95
IVEKANPVND
HA1 87-95
IVEK...PGDF
HA1 87-102
IVEK...DFND
HA1 87-104
KANP...PGDF
HA1 90-102
KANP...DFND
HA1 90-104
LCYPGDFND
HA1 96-104
LCYPGDFNDY
HA1 96-105
LCYPGDF
HA1 96-102
NDYE...INHF
HA1 103-118
YEEL...INHF
HA1 105-118
EELK...INHF
HA1 106-118
EKIQ...SSHE
HA1 119-131
EKIQ...EASL
HA1 119-133a
GVSS...KSSF
HA1 134-147
FRNVVWL
HA1 148-154
FRNVVW
HA1 148-153
LVMW...QTKL
HA1 177-194
LVMW...DAAE
HA1 177-190
VMWG...QTKL
HA1 178-194
VMWG...DAAE
HA1 178-190
WGIH...QTKL
HA1 180-194
YQNPTTY
HA1 195-201
YQNP...TSTL
HA1 195-209
YISVGTSTL
HA1 201-209
ISVGTSTL
HA1 202-209
NQRLVPRIAT
HA1 210-219
NQRL...RMEF**
HA1 210-232
RSKV...RMEF**
HA1 220-232
FWTILKPND
HA1 233-241
FWTI...AINF
HA1 233-245
WTIL
HA1 234-237
WTIL...AINF
HA1 234-245
WTILKPND
HA1 234-241
AINF
HA1 242-245
ESNGNFIAPEY
HA1 246-256
ESNGNFIAPEYA
HA1 246-257
IAPE...STIM
HA1 252-268
AYKI...STIM
HA1 257-268
AYKIVKKGDST
HA1 257-266
% Exchange
0
25
50
75
100
10
$10^2$
$10^3$
$10^4$
Time (seconds)
WT pH 7.4
WT pH 6.1
Mutant pH 7.4
Mutant pH 6.1
Mutant pH 5.9

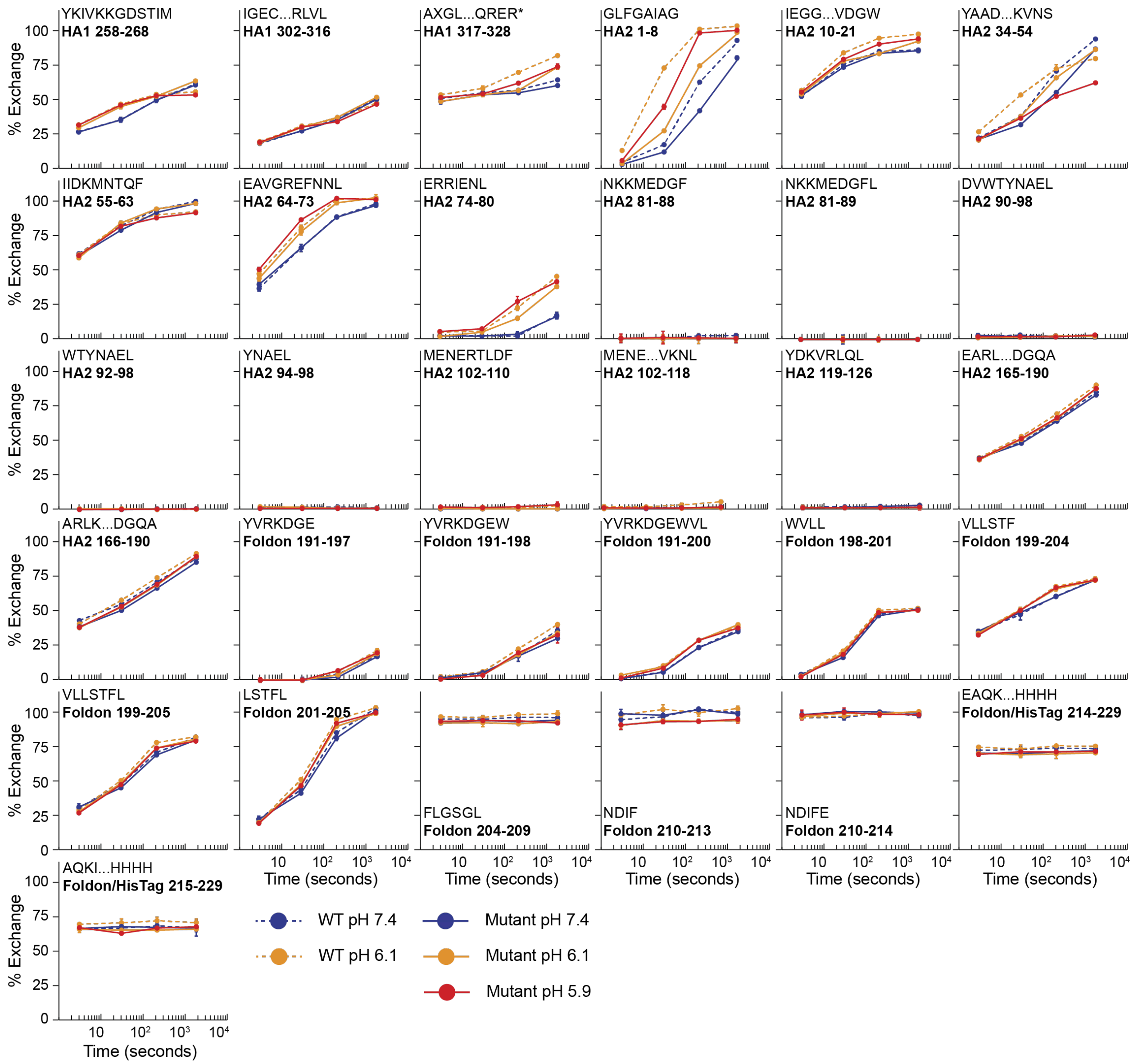


**Figure S2: Deuterium uptake plots for Viet04 WT and mutant**

Deuterium uptake plots for WT (dashed lines) and mutant (solid lines) at pH 7.4 (blue), 6.1 (orange), and 5.9 (red). Each timepoint is the average of at least two replicate measurements and error bars show the standard deviation between replicates. The data was corrected for in-exchange using a zero-second control. Percent exchange was determined using a totally deuterated control. Asterisks denote sequences that contain mutations.

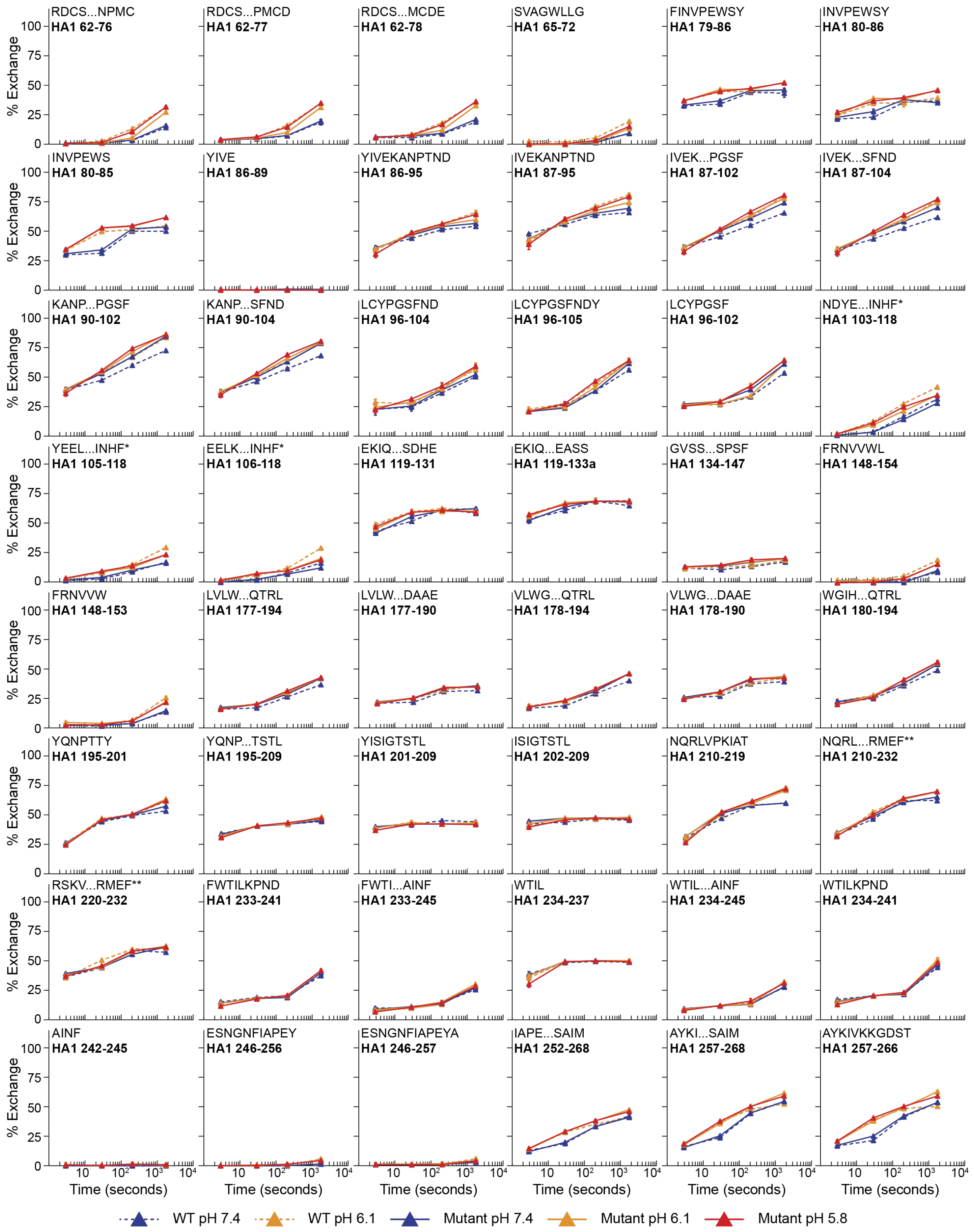
RDCS...NPMC HA1 62-76
RDCS...PMCD HA1 62-77
RDCS...MCDE HA1 62-78
SVAGWLLG HA1 65-72
FINVPEWSY HA1 79-86
INVPEWSY HA1 80-86
INVPEWS HA1 80-85
YIVE HA1 86-89
YIVEKANPTND HA1 86-95
IVEKANPTND HA1 87-95
IVEK...PGSF HA1 87-102
IVEK...SFND HA1 87-104
KANP...PGSF HA1 90-102
KANP...SFND HA1 90-104
LCYPGSFND HA1 96-104
LCYPGSFNDY HA1 96-105
LCYPGSF HA1 96-102
NDYE...INHF* HA1 103-118
YEEL...INHF* HA1 105-118
EELK...INHF* HA1 106-118
EKIQ...SDHE HA1 119-131
EKIQ...EASS HA1 119-133a
GVSS...SPSF HA1 134-147
FRNVVWL HA1 148-154
FRNVVW HA1 148-153
LVLW...QTRL HA1 177-194
LVLW...DAAE HA1 177-190
VLWG...QTRL HA1 178-194
VLWG...DAAE HA1 178-190
WGIH...QTRL HA1 180-194
YQNPTTY HA1 195-201
YQNP...TSTL HA1 195-209
YISIGTSTL HA1 201-209
ISIGTSTL HA1 202-209
NQRLVPKIAT HA1 210-219
NQRL...RMEF** HA1 210-232
RSKV...RMEF** HA1 220-232
FWTILKPND HA1 233-241
FWTI...AINF HA1 233-245
WTIL HA1 234-237
WTIL...AINF HA1 234-245
WTILKPND HA1 234-241
AINF HA1 242-245
ESNGNFIAPEY HA1 246-256
ESNGNFIAPEYA HA1 246-257
IAPE...SAIM HA1 252-268
AYKI...SAIM HA1 257-268
AYKIVKKGDST HA1 257-266
% Exchange
0
25
50
75
100
10
$10^2$
$10^3$
$10^4$
Time (seconds)
WT pH 7.4
WT pH 6.1
Mutant pH 7.4
Mutant pH 6.1
Mutant pH 5.8

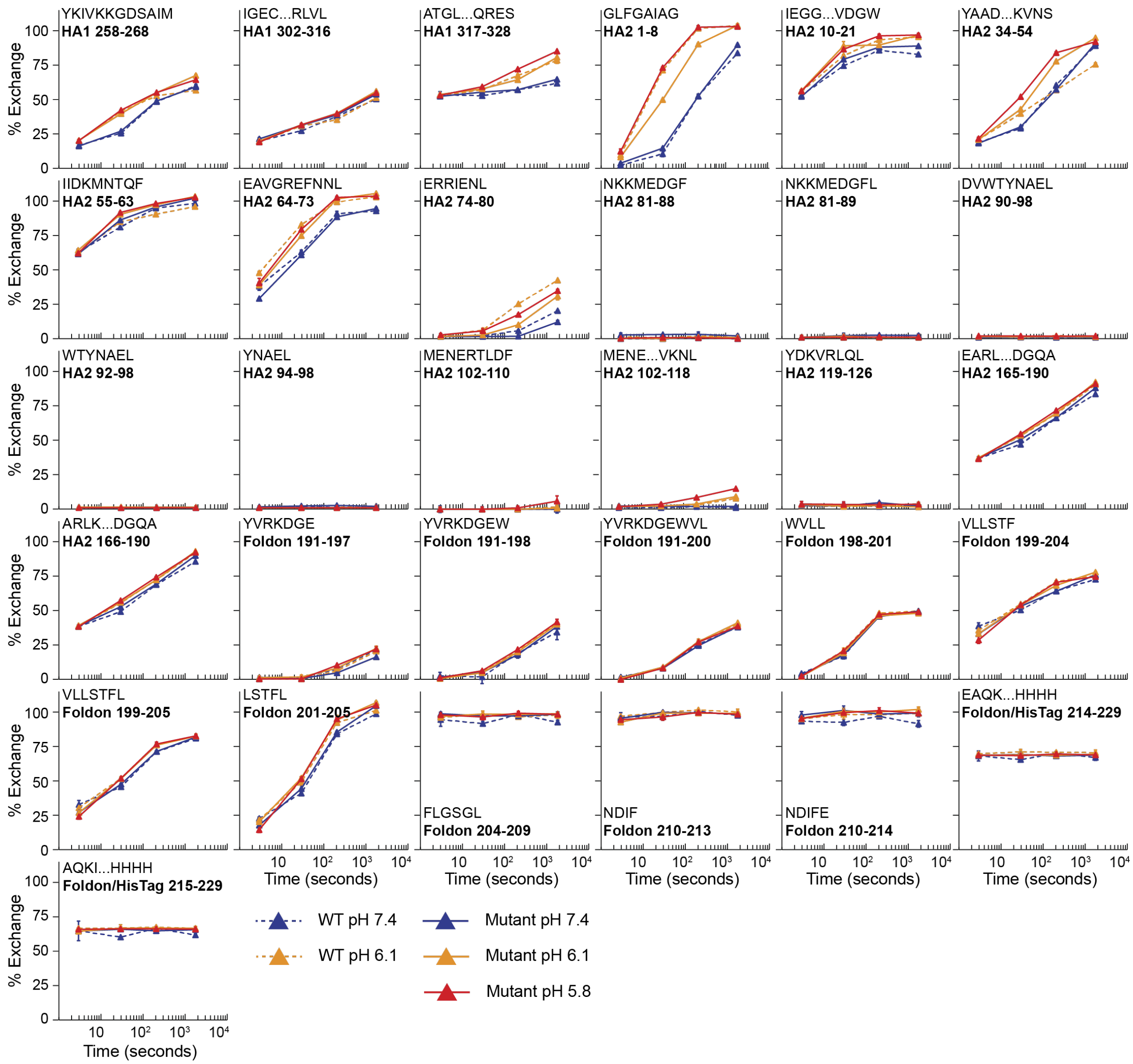


**Figure S3: Deuterium uptake plots for Indo05 WT and mutant**
Deuterium uptake plots for WT (dashed lines) and mutant (solid lines) at pH 7.4 (blue), 6.1 (orange), and 5.8 (red). Each timepoint is the average of at least two replicate measurements and error bars show the standard deviation between replicates. The data was corrected for in-exchange using a zero-second control. Percent exchange was determined using a totally deuterated control. Asterisks denote sequences that contain mutations.

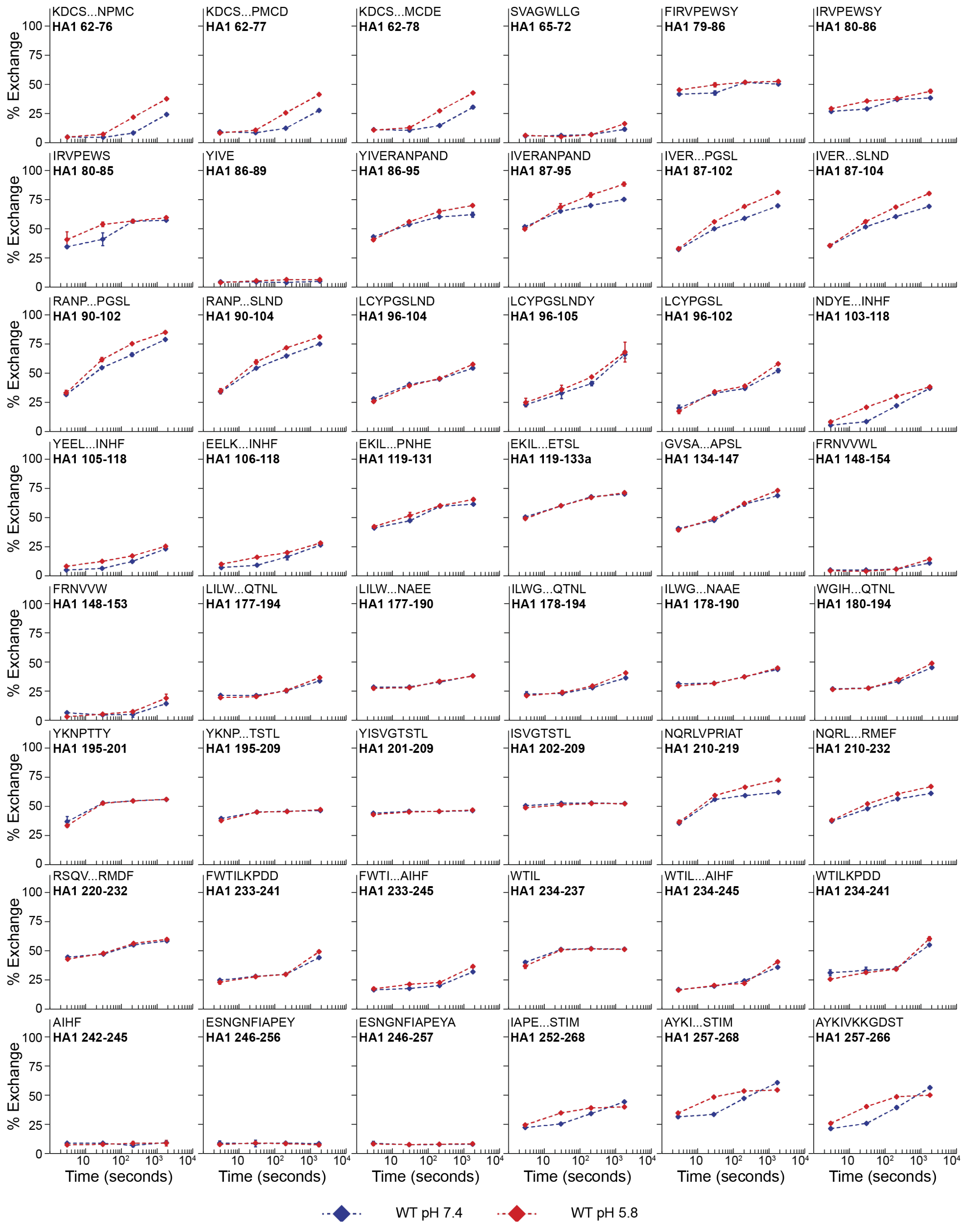

KDCS...NPMC
HA1 62-76
KDCS...PMCD
HA1 62-77
KDCS...MCDE
HA1 62-78
SVAGWLLG
HA1 65-72
FIRVPEWSY
HA1 79-86
IRVPEWSY
HA1 80-86
IRVPEWS
HA1 80-85
YIVE
HA1 86-89
YIVERANPAND
HA1 86-95
IVERANPAND
HA1 87-95
IVER...PGSL
HA1 87-102
IVER...SLND
HA1 87-104
RANP...PGSL
HA1 90-102
RANP...SLND
HA1 90-104
LCYPGSLND
HA1 96-104
LCYPGSLNDY
HA1 96-105
LCYPGSL
HA1 96-102
NDYE...INHF
HA1 103-118
YEEL...INHF
HA1 105-118
EELK...INHF
HA1 106-118
EKIL...PNHE
HA1 119-131
EKIL...ETSL
HA1 119-133a
GVSA...APSL
HA1 134-147
FRNVVWL
HA1 148-154
FRNVVW
HA1 148-153
LILW...QTNL
HA1 177-194
LILW...NAEE
HA1 177-190
ILWG...QTNL
HA1 178-194
ILWG...NAAE
HA1 178-190
WGIH...QTNL
HA1 180-194
YKNPTTY
HA1 195-201
YKNP...TSTL
HA1 195-209
YISVGTSTL
HA1 201-209
ISVGTSTL
HA1 202-209
NQRLVPRIAT
HA1 210-219
NQRL...RMEF
HA1 210-232
RSQV...RMDF
HA1 220-232
FWTILKPDD
HA1 233-241
FWTI...AIHF
HA1 233-245
WTIL
HA1 234-237
WTIL...AIHF
HA1 234-245
WTILKPDD
HA1 234-241
AIHF
HA1 242-245
ESNGNFIAPEY
HA1 246-256
ESNGNFIAPEYA
HA1 246-257
IAPE...STIM
HA1 252-268
AYKI...STIM
HA1 257-268
AYKIVKKGDST
HA1 257-266
% Exchange
0
25
50
75
100
10
$10^2$
$10^3$
$10^4$
Time (seconds)
WT pH 7.4
WT pH 5.8

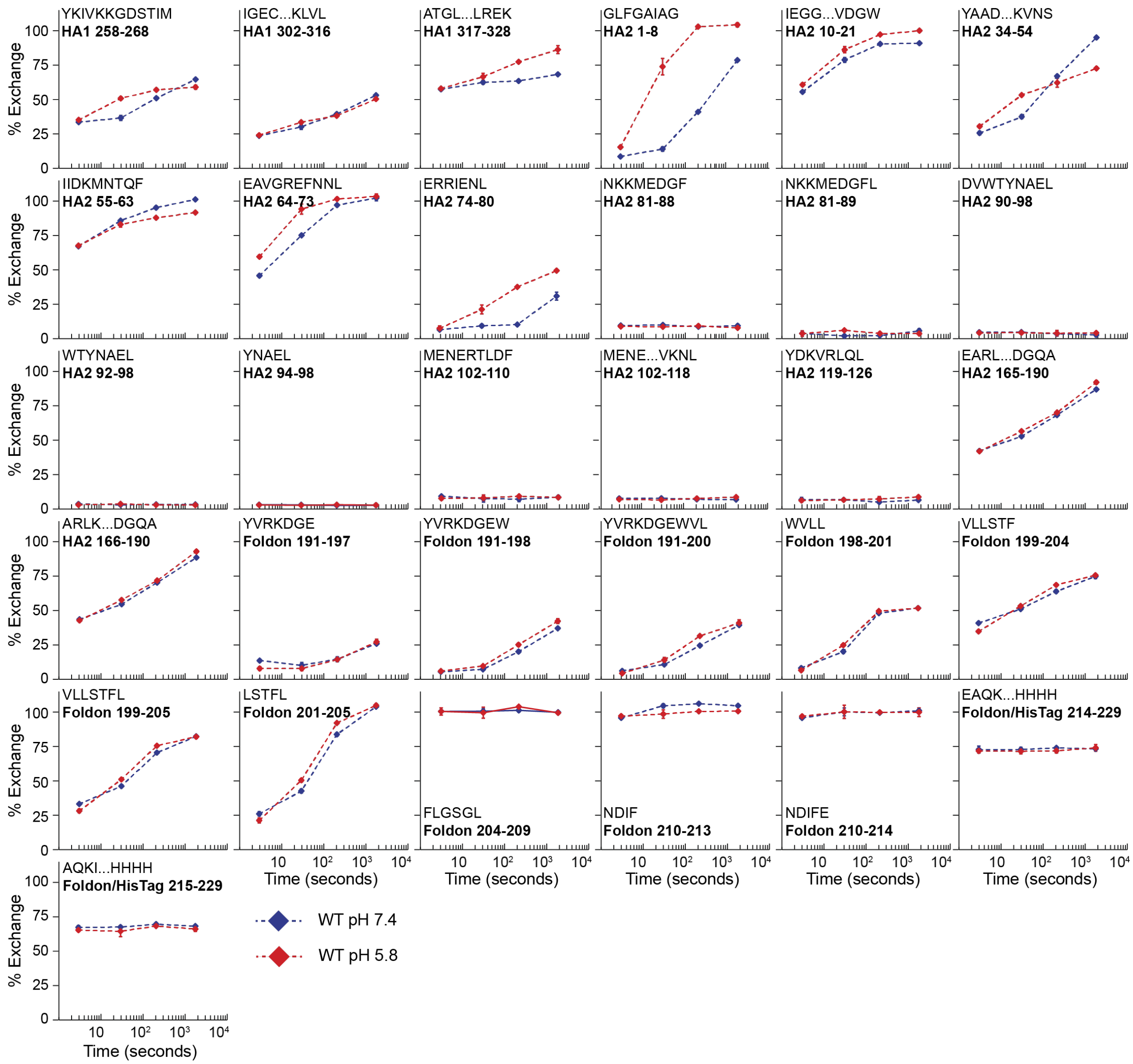


**Figure S4: Deuterium uptake plots for Colo22**

Deuterium uptake plots for Colo22 at pH 7.4 (blue) and 5.8 (red). Each timepoint is the average of at least two replicate measurements and error bars show the standard deviation between replicates. Percent exchange was determined using a totally deuterated control.

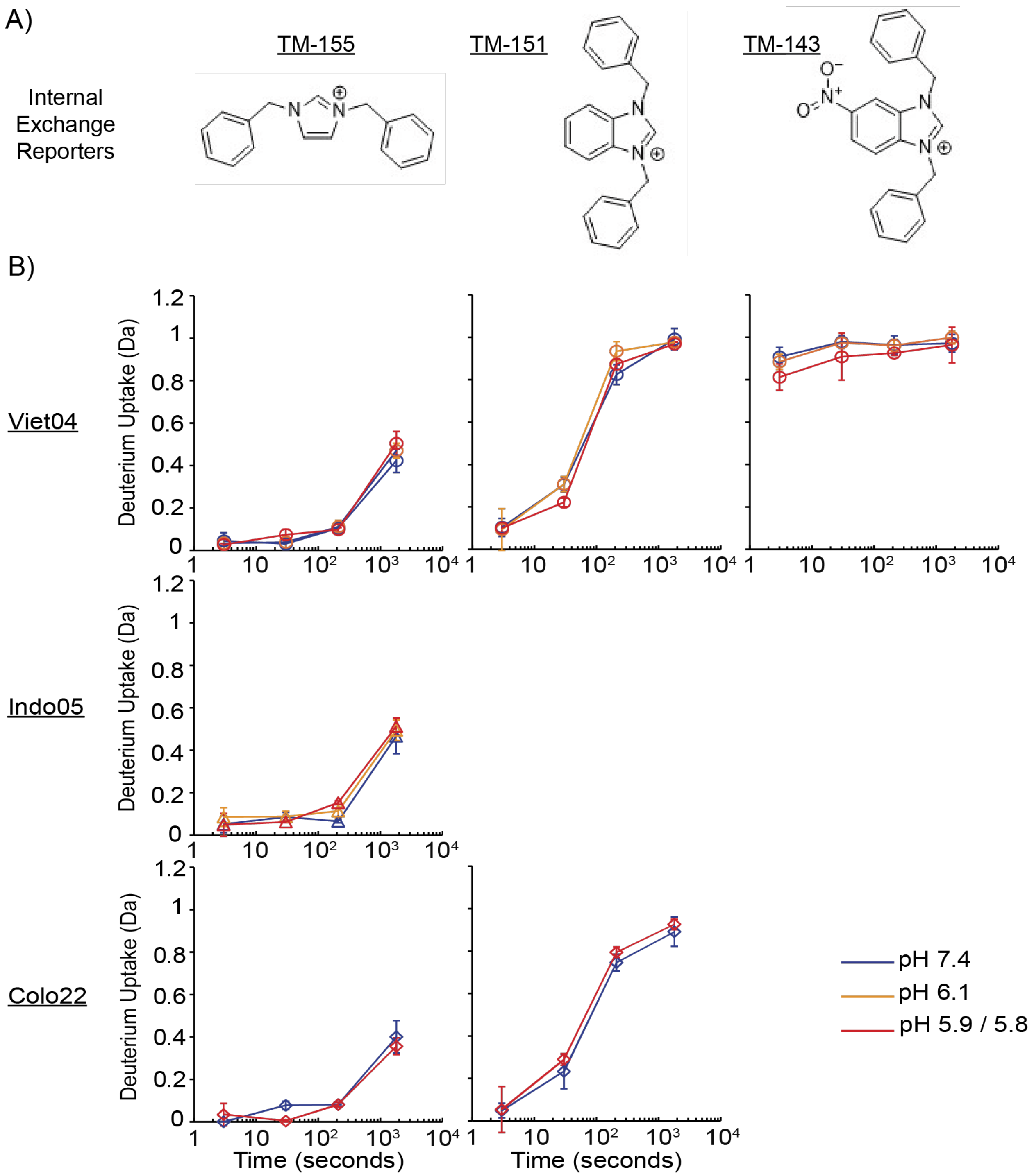


**Figure S5: Internal exchange reporters for HDX-MS**

**(A)** Structures of 3 internal exchange reporters (IER) present in HDX reactions and the corresponding deuterium uptake plots **(B)**. Due to low signal intensity in some samples, not all of the IERs could be monitored in every dataset. Exchange at each timepoint is the average of replicate measurements (n=4,3,2). Error bars are the standard deviation. For pH 7.4 and 6.1, the WT and mutant data was combined.

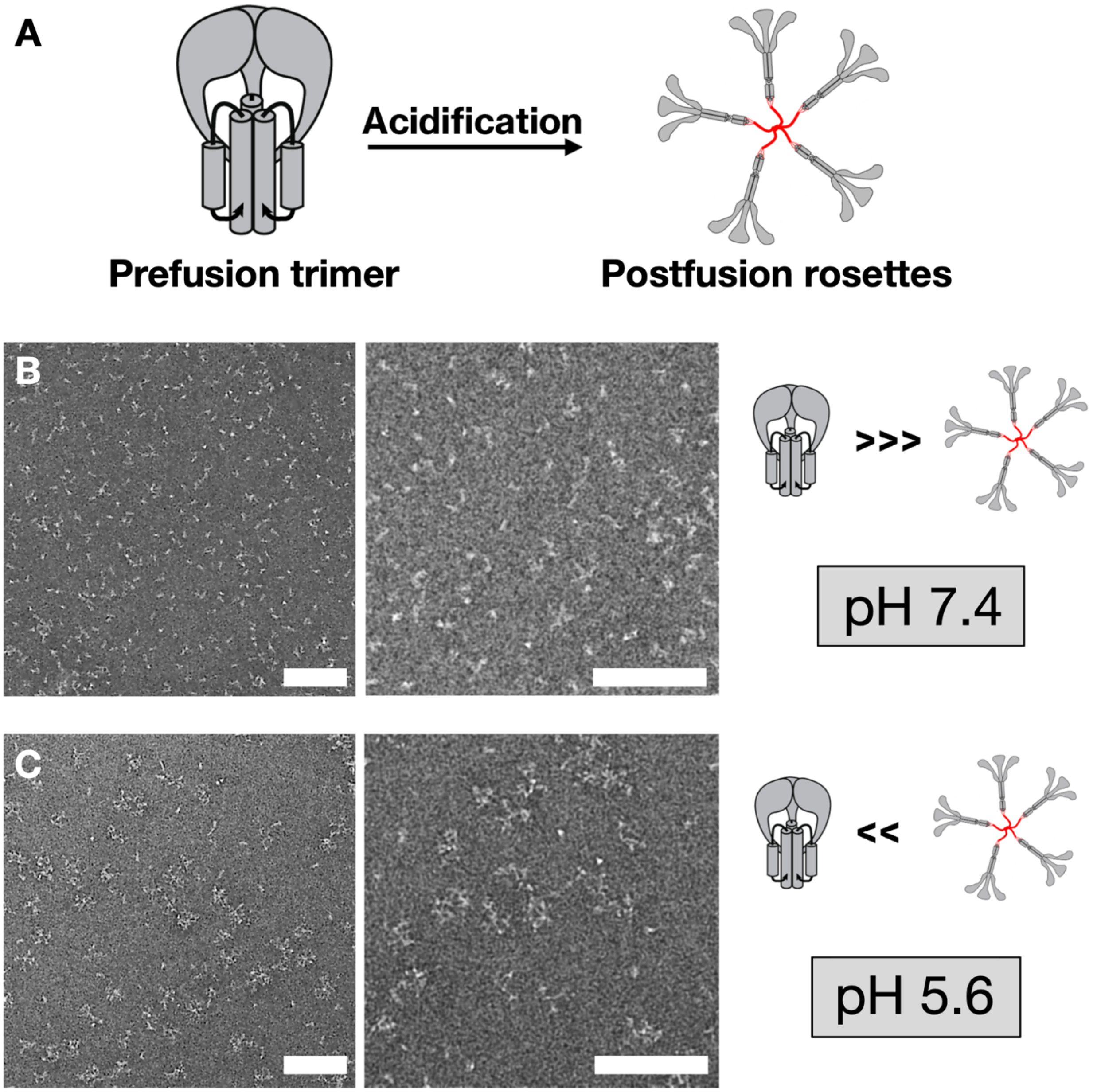


**Figure S6: Exposing HA to pH below the threshold of activation leads to formation of rosettes consistent with postfusion HA associating via hydrophobic fusion peptides.**
**(A)** Cartoon illustrating prefusion Colo22 HA monodisperse trimer and Colo22 HA as rosettes containing several postfusion trimers. **(B,C)** Negative stain electron micrographs after 1 hour of incubation at pH 7.4 and 5.6 respectively, indicating a large increase in postfusion rosette abundance below the threshold for activation. These provide an indication of the nature of the HA aggregate that is retained in BN-PAGE wells. Scale bars correspond to 100 nm.

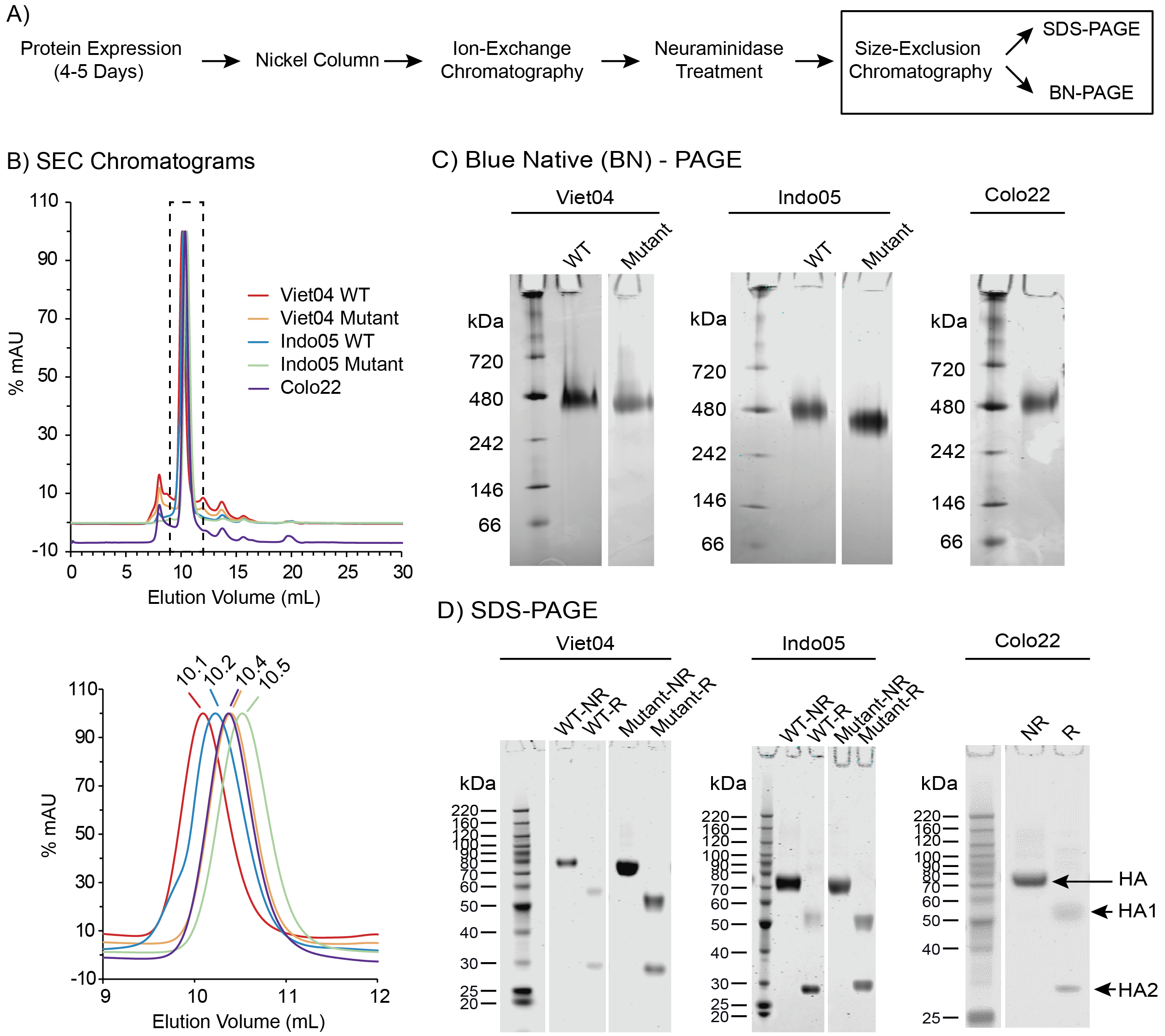


**Figure S7: Purification and characterization recombinant H5 HA**
**(A)** HA purification workflow. **(B)** Overlayed SEC chromatograms for all H5 constructs. The absorbance intensity was normalized to the highest signal for each protein. HA eluted as a single peak with retention times between 10.1-10.5 minutes. To assess trimer purity, each HA was characterized by BN-PAGE **(C)** and SDS-PAGE **(D)**. SDS-PAGE was performed under reducing (R) and nonreducing (NR) conditions. The lack of full-length HA under reducing condition suggests that there no uncleaved HA0 in our sample.

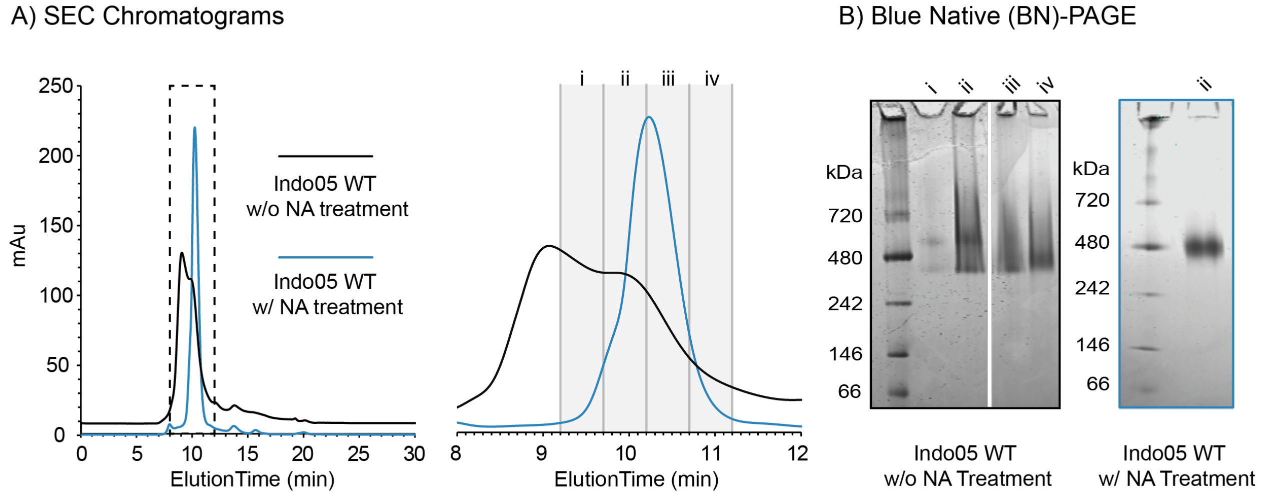


**Figure S8: Neuraminidase (NA) treatment**
**(A)** Overlayed SEC chromatograms for untreated (black) and treated (blue) Indo05 WT HA. Without NA-treatment, HA elutes as two unresolved peaks. Treating HA with NA prior to SEC, yields a single peak with a slightly shifted retention time. **(B)** BN-PAGE of individual fractions shows multiple MW species present in untreated HA. A single band corresponding to trimeric HA is seen in the treated sampl

| Peptide | Start | End | Sum %Exchange across timepoints | | | | | Differences | | | | |
|---|---|---|---|---|---|---|---|---|---|---|---|---|
| | | | WT pH 7.4 | WT pH 6.1 | Mutant pH 7.4 | Mutant pH 6.1 | Mutant pH 5.8 | WT/Mutant pH 7.4 | WT/Mutant pH 6.1/5.8 | WT pH 7.4/6.1 | Mutant pH 7.4/6.1 | Mutant pH 7.4/5.8 |
| RDCSVAGWLLGNPMC | 62 | 76 | 15 | 44 | 23 | 41 | 54 | 7 | 10 | 29 | 18 | 31 |
| RDCSVAGWLLGNPMCD | 62 | 77 | 29 | 56 | 36 | 52 | 68 | 7 | 11 | 27 | 16 | 32 |
| RDCSVAGWLLGNPMCDE | 62 | 78 | 34 | 63 | 41 | 59 | 71 | 7 | 9 | 29 | 18 | 30 |
| SVAGWLLG | 65 | 72 | 0 | 10 | 7 | 12 | 18 | 7 | 7 | 10 | 6 | 11 |
| FINVPEWSY | 79 | 86 | 155 | 177 | 154 | 184 | 175 | -1 | -2 | 22 | 29 | 20 |
| INVPEWSY | 80 | 86 | 113 | 146 | 122 | 146 | 148 | 9 | 2 | 33 | 24 | 26 |
| INVPEWS | 80 | 85 | 153 | 183 | 153 | 203 | 192 | 0 | 9 | 30 | 50 | 39 |
| YIVE | 86 | 89 | 2 | 0 | 1 | 0 | 5 | -1 | 5 | -2 | -1 | 4 |
| YIVEKANPVND | 86 | 95 | 169 | 191 | 180 | 181 | 193 | 11 | 2 | 22 | 1 | 13 |
| IVEKANPVND | 87 | 95 | 201 | 232 | 211 | 224 | 230 | 10 | -2 | 31 | 13 | 20 |
| IVEKANPVNDLCYPGDF | 87 | 102 | 171 | 200 | 200 | 209 | 223 | 28 | 23 | 29 | 9 | 23 |
| IVEKANPVNDLCYPGDFND | 87 | 104 | 168 | 194 | 193 | 203 | 215 | 25 | 20 | 26 | 10 | 22 |
| KANPVNDLCYPGDF | 90 | 102 | 181 | 210 | 218 | 228 | 242 | 37 | 32 | 29 | 10 | 23 |
| KANPVNDLCYPGDFND | 90 | 104 | 175 | 201 | 207 | 216 | 231 | 32 | 30 | 26 | 9 | 24 |
| LCYPGDFND | 96 | 104 | 104 | 111 | 137 | 140 | 145 | 33 | 34 | 7 | 4 | 8 |
| LCYPGDFNDY | 96 | 105 | 108 | 120 | 128 | 145 | 150 | 20 | 30 | 12 | 16 | 22 |
| LCYPGDF | 96 | 102 | 108 | 114 | 146 | 161 | 162 | 38 | 47 | 7 | 15 | 16 |
| NDYEELKHLLSRINHF | 103 | 118 | 52 | 80 | 47 | 78 | 80 | -5 | -1 | 28 | 30 | 32 |
| YEELKHLLSRINHF | 105 | 118 | 31 | 47 | 29 | 51 | 50 | -2 | 2 | 16 | 22 | 21 |
| EELKHLLSRINHF | 106 | 118 | 33 | 50 | 32 | 49 | 53 | -2 | 3 | 17 | 18 | 21 |
| EKIQIIPKSSWSSHE | 119 | 131 | 229 | 240 | 232 | 232 | 232 | 3 | -8 | 11 | 0 | 0 |
| EKIQIIPKSSWSSHEASL | 119 | 133a | 261 | 269 | 259 | 256 | 260 | -2 | -10 | 8 | -4 | 0 |
| GVSSACPYQGKSSF | 134 | 147 | 262 | 280 | 312 | 312 | 318 | 50 | 38 | 18 | 0 | 5 |
| FRNVVWL | 148 | 154 | 7 | 12 | 6 | 12 | 16 | -1 | 4 | 5 | 6 | 10 |
| FRNVVW | 148 | 153 | 12 | 11 | 6 | 16 | 24 | -6 | 12 | 0 | 10 | 18 |
| LVMWGIHHPNDAAEQTKL | 177 | 194 | 101 | 106 | 102 | 110 | 111 | 1 | 5 | 5 | 9 | 10 |
| LVMWGIHHPNDAAE | 177 | 190 | 117 | 117 | 115 | 114 | 116 | -2 | -1 | 0 | -2 | 1 |
| VMWGIHHPNDAAEQTKL | 178 | 194 | 110 | 120 | 109 | 115 | 122 | -1 | 2 | 10 | 6 | 13 |
| VMWGIHHPNDAAE | 178 | 190 | 132 | 137 | 133 | 132 | 132 | 1 | -4 | 4 | -1 | -1 |
| WGIHHPNDAAEQTKL | 180 | 194 | 127 | 140 | 136 | 142 | 156 | 9 | 16 | 13 | 6 | 20 |
| YQNPTTY | 195 | 201 | 184 | 190 | 185 | 194 | 199 | 1 | 9 | 6 | 9 | 14 |
| YQNPTTYISVGTSTL | 195 | 209 | 158 | 160 | 159 | 160 | 160 | 1 | 0 | 2 | 1 | 1 |

| | | | | | | | | | | | | |
|---|---|---|---|---|---|---|---|---|---|---|---|---|
| YISVGTSTL | 201 | 209 | 150 | 147 | 147 | 153 | 149 | -3 | 2 | -3 | 6 | 2 |
| ISVGTSTL | 202 | 209 | 179 | 174 | 176 | 174 | 176 | -3 | 1 | -5 | -2 | 0 |
| NQRLVPRIAT | 210 | 219 | 184 | 199 | 208 | 227 | 232 | 24 | 34 | 15 | 20 | 25 |
| NQRLVPRIATRSKVNGQSGRMEF | 210 | 232 | 190 | 201 | 214 | 223 | 227 | 24 | 26 | 11 | 9 | 13 |
| RSKVNGQSGRMEF | 220 | 232 | 189 | 196 | 208 | 213 | 221 | 20 | 24 | 8 | 4 | 12 |
| FWTILKPND | 233 | 241 | 90 | 91 | 87 | 90 | 95 | -3 | 4 | 1 | 3 | 8 |
| FWTILKPNDAINF | 233 | 245 | 57 | 60 | 52 | 59 | 58 | -6 | -1 | 2 | 7 | 7 |
| WTIL | 234 | 237 | 184 | 182 | 196 | 189 | 188 | 12 | 6 | -2 | -8 | -8 |
| WTILKPNDAINF | 234 | 245 | 63 | 65 | 58 | 59 | 65 | -4 | 0 | 2 | 0 | 7 |
| WTILKPND | 234 | 241 | 101 | 108 | 95 | 107 | 107 | -6 | -1 | 7 | 12 | 12 |
| AINF | 242 | 245 | 0 | 0 | 1 | 2 | 0 | 1 | 0 | 0 | 1 | -1 |
| ESNGNFIAPEY | 246 | 256 | 4 | 9 | 0 | 2 | 5 | -4 | -4 | 5 | 2 | 5 |
| ESNGNFIAPEYA | 246 | 257 | 4 | 8 | 0 | 4 | 6 | -4 | -2 | 4 | 4 | 6 |
| IAPEYAYKIVKKGDSTIM | 252 | 268 | 118 | 128 | 115 | 137 | 128 | -3 | 0 | 10 | 23 | 13 |
| AYKIVKKGDSTIM | 257 | 268 | 158 | 169 | 153 | 172 | 166 | -5 | -3 | 11 | 19 | 12 |
| AYKIVKKGDST | 257 | 266 | 135 | 155 | 141 | 167 | 159 | 6 | 3 | 21 | 26 | 18 |
| YKIVKKGDSTIM | 258 | 268 | 171 | 185 | 169 | 188 | 181 | -1 | -4 | 15 | 18 | 12 |
| IGECPKYVKSNRLVL | 302 | 316 | 129 | 131 | 129 | 134 | 127 | -1 | -4 | 1 | 5 | -2 |
| ATGLRNSPQRER | 317 | 328 | 227 | 263 | 217 | 233 | 242 | -10 | -22 | 36 | 15 | 24 |
| GLFGAIAG | 1 | 8 | 177 | 291 | 137 | 204 | 249 | -40 | -41 | 113 | 67 | 112 |
| IEGGWQGMVDGW | 10 | 21 | 299 | 333 | 296 | 308 | 319 | -3 | -14 | 34 | 12 | 24 |
| YAADKESTQKAIDGVTNKVNS | 34 | 54 | 222 | 229 | 192 | 208 | 170 | -30 | -60 | 7 | 16 | -22 |
| IIDKMNTQF | 55 | 63 | 337 | 326 | 328 | 334 | 320 | -8 | -5 | -11 | 6 | -8 |
| EAVGREFNNL | 64 | 73 | 289 | 332 | 291 | 323 | 340 | 2 | 8 | 42 | 32 | 49 |
| ERRIENL | 74 | 80 | 21 | 76 | 22 | 57 | 79 | 0 | 3 | 55 | 36 | 57 |
| NKKMEDGF | 81 | 88 | 2 | 0 | 0 | 0 | 2 | -2 | 2 | -2 | 0 | 2 |
| NKKMEDGFL | 81 | 89 | 0 | 0 | 0 | 0 | 0 | 0 | 0 | 0 | 0 | 0 |
| DVWTYNAEL | 90 | 98 | 7 | 6 | 6 | 4 | 6 | -1 | 0 | -1 | -2 | -1 |
| WTYNAEL | 92 | 98 | 1 | 0 | 1 | 2 | 1 | -1 | 0 | -1 | 1 | 0 |
| YNAEL | 94 | 98 | 3 | 0 | 0 | 2 | 0 | -3 | 0 | -3 | 2 | 0 |
| MENERTLDF | 102 | 110 | 0 | 5 | 0 | 0 | 7 | 0 | 2 | 5 | 0 | 7 |
| MENERTLDFHDSNVKNL | 102 | 118 | 3 | 10 | 2 | 1 | 3 | -2 | -8 | 7 | 0 | 1 |
| YDKVRLQL | 119 | 126 | 2 | 3 | 2 | 0 | 0 | 0 | -3 | 1 | -2 | -2 |
| EARLKREEISGVGSGYIPEAPRDGQA | 165 | 190 | 234 | 248 | 230 | 238 | 240 | -4 | -8 | 14 | 8 | 10 |
| ARLKREEISGVGSGYIPEAPRDGQA | 166 | 190 | 253 | 261 | 237 | 245 | 247 | -16 | -14 | 7 | 8 | 10 |
| YVRKDGE | 191 | 197 | 22 | 26 | 17 | 21 | 25 | -5 | 0 | 3 | 4 | 9 |

| | | | | | | | | | | | | |
|---|---|---|---|---|---|---|---|---|---|---|---|---|
| YVRKDGEW | 191 | 198 | 57 | 68 | 52 | 52 | 53 | -5 | -15 | 11 | 1 | 1 |
| YVRKDGEWVL | 191 | 200 | 62 | 75 | 61 | 78 | 73 | -2 | -2 | 12 | 17 | 12 |
| WVLL | 198 | 201 | 119 | 124 | 115 | 119 | 118 | -4 | -6 | 5 | 4 | 3 |
| VLLSTF | 199 | 204 | 217 | 228 | 218 | 224 | 224 | 1 | -5 | 11 | 6 | 5 |
| VLLSTFL | 199 | 205 | 232 | 240 | 226 | 231 | 228 | -6 | -12 | 8 | 6 | 2 |
| LSTFL | 201 | 205 | 252 | 267 | 244 | 254 | 256 | -9 | -12 | 15 | 10 | 12 |
| FLGSGL | 204 | 209 | 384 | 391 | 375 | 370 | 374 | -9 | -17 | 7 | -4 | 0 |
| NDIF | 210 | 213 | 393 | 401 | 396 | 371 | 371 | 4 | -30 | 9 | -25 | -25 |
| NDIFE | 210 | 214 | 389 | 391 | 398 | 395 | 395 | 9 | 4 | 2 | -3 | -4 |
| EAQKIEWHEGHHHHHH | 214 | 229 | 292 | 298 | 283 | 279 | 283 | -9 | -15 | 6 | -4 | 0 |
| AQKIEWHEGHHHHHH | 215 | 229 | 266 | 280 | 266 | 260 | 262 | 0 | -19 | 15 | -6 | -4 |

**Table S1: Differences in total exchange (Viet04), related to heatmaps**
For each peptide, the total percent exchange was determined by adding the percent exchange at all timepoints. The difference in total exchange is shown for comparisons that are discussed in the main text. Differences are colored to match how they appear in the related heatmap figures.

| Peptide | Start | End | Sum %Exchange across timepoints | | | | | Differences | | | | |
|---|---|---|---|---|---|---|---|---|---|---|---|---|
| | | | WT pH 7.4 | WT pH 6.1 | Mutant pH 7.4 | Mutant pH 6.1 | Mutant pH 5.8 | WT/Mutant pH 7.4 | WT/Mutant pH 6.1/5.8 | WT pH 7.4/6.1 | Mutant pH 7.4/6.1 | Mutant pH 7.4/5.8 |
| RDCSVAGWLLGNPMC | 62 | 76 | 17 | 46 | 19 | 32 | 42 | 2 | -4 | 29 | 13 | 23 |
| RDCSVAGWLLGNPMCD | 62 | 77 | 34 | 60 | 35 | 50 | 60 | 1 | 0 | 25 | 15 | 24 |
| RDCSVAGWLLGNPMCDE | 62 | 78 | 39 | 68 | 44 | 58 | 66 | 5 | -2 | 29 | 14 | 23 |
| SVAGWLLG | 65 | 72 | 14 | 31 | 13 | 17 | 21 | -1 | -11 | 17 | 4 | 8 |
| FINVPEWSY | 79 | 86 | 154 | 172 | 162 | 182 | 181 | 8 | 9 | 18 | 20 | 20 |
| INVPEWSY | 80 | 86 | 118 | 134 | 125 | 149 | 150 | 7 | 16 | 17 | 24 | 25 |
| INVPEWS | 80 | 85 | 161 | 189 | 171 | 203 | 204 | 9 | 15 | 28 | 32 | 33 |
| YIVE | 86 | 89 | 1 | 0 | 2 | 0 | 1 | 1 | 0 | -1 | -2 | -2 |
| YIVEKANPTND | 86 | 95 | 187 | 207 | 195 | 200 | 201 | 7 | -6 | 20 | 5 | 7 |
| IVEKANPTND | 87 | 95 | 235 | 255 | 238 | 245 | 251 | 3 | -4 | 20 | 7 | 12 |
| IVEKANPTNDLCYPGSF | 87 | 102 | 200 | 225 | 219 | 227 | 229 | 19 | 4 | 25 | 8 | 10 |
| IVEKANPTNDLCYPGSFND | 87 | 104 | 189 | 217 | 208 | 216 | 220 | 19 | 3 | 27 | 8 | 11 |
| KANPTNDLCYPGSF | 90 | 102 | 219 | 243 | 245 | 253 | 252 | 25 | 9 | 24 | 8 | 8 |
| KANPTNDLCYPGSFND | 90 | 104 | 209 | 231 | 229 | 235 | 238 | 20 | 6 | 22 | 5 | 9 |
| LCYPGSFND | 96 | 104 | 135 | 155 | 140 | 152 | 156 | 5 | 2 | 19 | 12 | 16 |
| LCYPGSFNDY | 96 | 105 | 141 | 157 | 144 | 151 | 158 | 3 | 1 | 16 | 7 | 14 |
| LCYPGSF | 96 | 102 | 141 | 149 | 158 | 164 | 162 | 17 | 13 | 8 | 6 | 4 |
| NDYEELKHLLSRINHF | 103 | 118 | 54 | 85 | 47 | 69 | 74 | -6 | -11 | 31 | 21 | 26 |
| YEELKHLLSRINHF | 105 | 118 | 27 | 52 | 29 | 44 | 47 | 2 | -6 | 26 | 15 | 18 |
| EELKHLLSRINHF | 106 | 118 | 27 | 49 | 23 | 40 | 39 | -4 | -11 | 22 | 17 | 16 |
| EKIQIIPKSSWSDHE | 119 | 131 | 215 | 233 | 220 | 224 | 226 | 5 | -7 | 18 | 5 | 6 |
| EKIQIIPKSSWSDHEASS | 119 | 133a | 247 | 261 | 252 | 258 | 259 | 5 | -2 | 14 | 6 | 7 |
| GVSSACPYLGSPSF | 134 | 147 | 50 | 55 | 61 | 62 | 64 | 11 | 9 | 5 | 0 | 3 |
| FRNVVWL | 148 | 154 | 11 | 29 | 10 | 22 | 18 | 0 | -11 | 18 | 12 | 8 |
| FRNVVW | 148 | 153 | 20 | 36 | 22 | 35 | 31 | 2 | -5 | 16 | 13 | 9 |
| LVLWGIHHPNDAAEQTRL | 177 | 194 | 96 | 108 | 108 | 109 | 110 | 12 | 3 | 11 | 1 | 2 |
| LVLWGIHHPNDAAE | 177 | 190 | 105 | 114 | 117 | 116 | 115 | 11 | 0 | 9 | 0 | -2 |
| VLWGIHHPNDAAEQTRL | 178 | 194 | 104 | 118 | 118 | 119 | 120 | 13 | 2 | 14 | 1 | 2 |
| VLWGIHHPNDAAE | 178 | 190 | 129 | 134 | 141 | 140 | 139 | 12 | 5 | 5 | -1 | -2 |
| WGIHHPNDAAEQTRL | 180 | 194 | 131 | 141 | 141 | 144 | 143 | 10 | 1 | 10 | 3 | 2 |
| YQNPTTY | 195 | 201 | 174 | 185 | 179 | 186 | 184 | 5 | -1 | 11 | 7 | 4 |
| YQNPTTYISIGTSTL | 195 | 209 | 162 | 162 | 162 | 164 | 162 | 0 | 0 | 0 | 2 | 0 |

| | | | | | | | | | | | |
|---|---|---|---|---|---|---|---|---|---|---|---|
| YISIGTSTL | 201 | 209 | 169 | 167 | 166 | 165 | 161 | -3 | -5 | -2 | 0 | -4 |
| ISIGTSTL | 202 | 209 | 178 | 180 | 187 | 185 | 180 | 9 | 0 | 2 | -2 | -7 |
| NQRLVPKIAT | 210 | 219 | 196 | 213 | 201 | 215 | 213 | 6 | 0 | 17 | 14 | 11 |
| NQRLVPKIATRSKVNGQSGRMEF | 210 | 232 | 204 | 218 | 210 | 218 | 217 | 6 | -2 | 14 | 8 | 7 |
| RSKVNGQSGRMEF | 220 | 232 | 197 | 207 | 201 | 203 | 203 | 4 | -4 | 10 | 2 | 2 |
| FWTILKPND | 233 | 241 | 92 | 94 | 91 | 94 | 91 | -1 | -2 | 2 | 3 | 1 |
| FWTILKPNDAINF | 233 | 245 | 60 | 59 | 60 | 64 | 60 | 0 | 1 | -1 | 4 | -1 |
| WTIL | 234 | 237 | 185 | 184 | 187 | 187 | 179 | 1 | -5 | -2 | 1 | -8 |
| WTILKPNDAINF | 234 | 245 | 62 | 66 | 62 | 67 | 67 | 0 | 1 | 3 | 4 | 4 |
| WTILKPND | 234 | 241 | 105 | 108 | 104 | 109 | 105 | 0 | -3 | 4 | 5 | 1 |
| AINF | 242 | 245 | 0 | 2 | 0 | 3 | 3 | 0 | 1 | 2 | 3 | 3 |
| ESNGNFIAPEY | 246 | 256 | 0 | 5 | 1 | 6 | 6 | 0 | 0 | 5 | 6 | 5 |
| ESNGNFIAPEYA | 246 | 257 | 2 | 5 | 2 | 5 | 2 | 0 | -3 | 3 | 3 | 0 |
| IAPEYAYKIVKKGDSAIM | 252 | 268 | 107 | 119 | 106 | 129 | 128 | -1 | 8 | 12 | 22 | 22 |
| AYKIVKKGDSAIM | 257 | 268 | 139 | 156 | 140 | 165 | 166 | 1 | 9 | 17 | 25 | 26 |
| AYKIVKKGDSA | 257 | 266 | 135 | 160 | 141 | 173 | 173 | 5 | 12 | 25 | 32 | 32 |
| YKIVKKGDSAIM | 258 | 268 | 151 | 171 | 151 | 182 | 182 | 1 | 11 | 20 | 31 | 31 |
| IGECPKYVKSNRLVL | 302 | 316 | 135 | 136 | 145 | 147 | 145 | 10 | 9 | 1 | 2 | 0 |
| ATGLRNSPQRES | 317 | 328 | 224 | 254 | 228 | 255 | 268 | 4 | 14 | 31 | 27 | 40 |
| GLFGAIAG | 1 | 8 | 150 | 287 | 161 | 253 | 292 | 11 | 5 | 137 | 92 | 131 |
| IEGGWQGMVDGW | 10 | 21 | 295 | 326 | 308 | 331 | 335 | 12 | 9 | 30 | 24 | 28 |
| YAADKESTQKAIDGVTNKVNS | 34 | 54 | 199 | 195 | 198 | 238 | 252 | -1 | 57 | -4 | 40 | 53 |
| IIDKMNTQF | 55 | 63 | 337 | 334 | 346 | 355 | 355 | 8 | 21 | -3 | 10 | 9 |
| EAVGREFNNL | 64 | 73 | 285 | 333 | 273 | 321 | 326 | -12 | -7 | 48 | 48 | 53 |
| ERRIENL | 74 | 80 | 29 | 75 | 15 | 44 | 59 | -14 | -16 | 46 | 28 | 44 |
| NKKMEDGF | 81 | 88 | 0 | 6 | 11 | 2 | 2 | 11 | -4 | 6 | -9 | -9 |
| NKKMEDGFL | 81 | 89 | 0 | 0 | 4 | 1 | 0 | 4 | 0 | 0 | -3 | -4 |
| DVWTYNAEL | 90 | 98 | 1 | 4 | 0 | 2 | 4 | -1 | 0 | 3 | 2 | 4 |
| WTYNAEL | 92 | 98 | 1 | 0 | 0 | 2 | 0 | -1 | 0 | -1 | 2 | 0 |
| YNAEL | 94 | 98 | 1 | 1 | 5 | 0 | 0 | 3 | -1 | 0 | -5 | -5 |
| MENERTLDF | 102 | 110 | 0 | 0 | 0 | 0 | 5 | 0 | 5 | 0 | 0 | 5 |
| MENERTLDFHDSNVKNL | 102 | 118 | 0 | 11 | 4 | 13 | 25 | 4 | 14 | 11 | 9 | 21 |
| YDKVRLQL | 119 | 126 | 8 | 8 | 10 | 8 | 12 | 2 | 4 | 0 | -2 | 2 |
| EARLKREEISGVGSGYIPEAPRDGQA | 165 | 190 | 233 | 249 | 242 | 253 | 254 | 8 | 4 | 16 | 11 | 12 |
| ARLKREEISGVGSGYIPEAPRDGQA | 166 | 190 | 242 | 258 | 251 | 260 | 262 | 9 | 4 | 16 | 10 | 11 |

| | | | | | | | | | | | |
|---|---|---|---|---|---|---|---|---|---|---|---|
| YVRKDGE | 191 | 197 | 31 | 25 | 20 | 32 | 30 | -11 | 5 | -6 | 12 | 10 |
| YVRKDGEW | 191 | 198 | 57 | 68 | 63 | 65 | 70 | 5 | 3 | 11 | 2 | 8 |
| YVRKDGEWVL | 191 | 200 | 73 | 76 | 73 | 79 | 73 | -1 | -3 | 3 | 6 | 0 |
| WVLL | 198 | 201 | 117 | 121 | 116 | 115 | 119 | -1 | -2 | 4 | -1 | 3 |
| VLLSTF | 199 | 204 | 224 | 235 | 224 | 231 | 226 | 0 | -9 | 11 | 7 | 2 |
| VLLSTFL | 199 | 205 | 227 | 237 | 224 | 233 | 232 | -3 | -6 | 11 | 9 | 8 |
| LSTFL | 201 | 205 | 246 | 264 | 252 | 273 | 266 | 6 | 2 | 18 | 21 | 14 |
| FLGSGL | 204 | 209 | 377 | 389 | 392 | 392 | 392 | 15 | 3 | 12 | 1 | 0 |
| NDIF | 210 | 213 | 392 | 398 | 394 | 390 | 389 | 2 | -10 | 7 | -3 | -5 |
| NDIFE | 210 | 214 | 376 | 392 | 398 | 400 | 397 | 23 | 5 | 16 | 1 | -2 |
| EAQKIEWHEGHHHHHH | 214 | 229 | 272 | 284 | 276 | 277 | 277 | 4 | -7 | 12 | 1 | 1 |
| AQKIEWHEGHHHHHH | 215 | 229 | 253 | 268 | 261 | 265 | 264 | 8 | -3 | 14 | 4 | 3 |

**Table S2: Differences in total exchange (Indo05), related to heatmaps**
For each peptide, the total percent exchange was determined by adding the percent exchange at all timepoints. The difference in total exchange is shown for comparisons that are discussed in the main text. Differences are colored to match how they appear in the related heatmap figures.

| Peptide | Start | End | Sum %Exchange across timepoints | | Differences |
|---|---|---|---|---|---|
| | | | WT pH 7.4 | WT pH 5.8 | WT pH7.4/5.8 |
| KDCSVAGWLLGNPMC | 62 | 76 | 40 | 70 | 30 |
| KDCSVAGWLLGNPMCD | 62 | 77 | 58 | 86 | 28 |
| KDCSVAGWLLGNPMCDE | 62 | 78 | 64 | 91 | 27 |
| SVAGWLLG | 65 | 72 | 29 | 33 | 4 |
| FIRVPEWSY | 79 | 86 | 187 | 200 | 13 |
| IRVPEWSY | 80 | 86 | 132 | 147 | 16 |
| IRVPEWS | 80 | 85 | 190 | 211 | 21 |
| YIVE | 86 | 89 | 16 | 20 | 4 |
| YIVERANPAND | 86 | 95 | 219 | 232 | 13 |
| IVERANPAND | 87 | 95 | 262 | 285 | 24 |
| IVERANPANDLCYPGSL | 87 | 102 | 213 | 241 | 28 |
| IVERANPANDLCYPGSLND | 87 | 104 | 215 | 239 | 24 |
| RANPANDLCYPGSL | 90 | 102 | 230 | 254 | 24 |
| RANPANDLCYPGSLND | 90 | 104 | 228 | 247 | 19 |
| LCYPGSLND | 96 | 104 | 168 | 168 | 0 |
| LCYPGSLNDY | 96 | 105 | 161 | 174 | 13 |
| LCYPGSL | 96 | 102 | 139 | 146 | 7 |
| NDYEELKHMLSRINHF | 103 | 118 | 72 | 96 | 24 |
| YEELKHMLSRINHF | 105 | 118 | 47 | 63 | 16 |
| EELKHMLSRINHF | 106 | 118 | 58 | 73 | 15 |
| EKILIIPKSSWPNHE | 119 | 131 | 207 | 217 | 10 |
| EKILIIPKSSWPNHETSL | 119 | 133a | 246 | 245 | -1 |
| GVSAACPYQGAPSF | 134 | 147 | 216 | 222 | 6 |
| FRNVVWL | 148 | 154 | 26 | 28 | 1 |
| FRNVVW | 148 | 153 | 29 | 33 | 4 |
| LILWGIHHSNNAEEQTNL | 177 | 194 | 100 | 101 | 0 |
| LILWGIHHSNNAEE | 177 | 190 | 127 | 126 | -1 |
| ILWGIHHSNNAEEQTNL | 178 | 194 | 105 | 110 | 5 |
| ILWGIHHSNNAEE | 178 | 190 | 143 | 142 | -1 |
| WGIHHSNNAEEQTNL | 180 | 194 | 132 | 136 | 5 |
| YKNPTTY | 195 | 201 | 198 | 194 | -4 |
| YKNPTTYISVGTSTL | 195 | 209 | 177 | 176 | -1 |
| YISVGTSTL | 201 | 209 | 181 | 180 | -1 |
| ISVGTSTL | 202 | 209 | 207 | 203 | -4 |

| | | | | | |
|---|---|---|---|---|---|
| NQRLAPKIAT | 210 | 219 | 214 | 236 | 22 |
| NQRLAPKIATRSQVNGQRGRMDF | 210 | 232 | 203 | 218 | 15 |
| RSQVNGQRGRMDF | 220 | 232 | 203 | 205 | 2 |
| FWTILKPDD | 233 | 241 | 125 | 128 | 3 |
| FWTILKPDDAIHF | 233 | 245 | 87 | 99 | 12 |
| WTIL | 234 | 237 | 192 | 188 | -4 |
| WTILKPDDAIHF | 234 | 245 | 94 | 97 | 3 |
| WTILKPDD | 234 | 241 | 151 | 148 | -3 |
| AIHF | 242 | 245 | 34 | 33 | -1 |
| ESNGNFIAPEY | 246 | 256 | 32 | 30 | -2 |
| ESNGNFIAPEYA | 246 | 257 | 29 | 30 | 0 |
| IAPEYAYKIVKKGDSTIM | 252 | 268 | 124 | 136 | 12 |
| AYKIVKKGDSTIM | 257 | 268 | 170 | 188 | 18 |
| AYKIVKKGDST | 257 | 266 | 144 | 166 | 22 |
| YKIVKKGDSTIM | 258 | 268 | 182 | 199 | 16 |
| IGECPKYVKSNKLVL | 302 | 316 | 145 | 145 | 0 |
| ATGLRNSPLREK | 317 | 328 | 248 | 285 | 37 |
| GLFGAIAG | 1 | 8 | 141 | 295 | 154 |
| IEGGWQGMVDGW | 10 | 21 | 315 | 344 | 29 |
| YAADKESTQKAIDGVTNKVNS | 34 | 54 | 224 | 217 | -7 |
| IIDKMNTQF | 55 | 63 | 348 | 328 | -19 |
| EAVGREFNNL | 64 | 73 | 319 | 357 | 38 |
| ERRIENL | 74 | 80 | 56 | 115 | 59 |
| NKKMEDGF | 81 | 88 | 36 | 33 | -3 |
| NKKMEDGFL | 81 | 89 | 15 | 18 | 3 |
| DVWTYNAEL | 90 | 98 | 16 | 17 | 1 |
| WTYNAEL | 92 | 98 | 10 | 9 | -1 |
| YNAEL | 94 | 98 | 12 | 12 | 0 |
| MENERTLDF | 102 | 110 | 30 | 31 | 1 |
| MENERTLDFHDSNVKNL | 102 | 118 | 31 | 32 | 0 |
| YDKVRLQL | 119 | 126 | 23 | 27 | 4 |
| EARLKREEISGVGSGYIPEAPRDGQA | 165 | 190 | 249 | 259 | 11 |
| ARLKREEISGVGSGYIPEAPRDGQA | 166 | 190 | 259 | 267 | 8 |
| YVRKDGE | 191 | 197 | 64 | 57 | -7 |
| YVRKDGEW | 191 | 198 | 71 | 84 | 13 |
| YVRKDGEWVL | 191 | 200 | 81 | 91 | 10 |

| | | | | | |
|---|---|---|---|---|---|
| WVLL | 198 | 201 | 124 | 129 | 5 |
| VLLSTF | 199 | 204 | 229 | 230 | 2 |
| VLLSTFL | 199 | 205 | 230 | 235 | 5 |
| LSTFL | 201 | 205 | 254 | 266 | 12 |
| FLGSGL | 204 | 209 | 403 | 404 | 1 |
| NDIF | 210 | 213 | 409 | 395 | -14 |
| NDIFE | 210 | 214 | 396 | 396 | 0 |
| EAQKIEWHEGHHHHHH | 214 | 229 | 291 | 287 | -4 |
| AQKIEWHEGHHHHHH | 215 | 229 | 273 | 264 | -9 |

**Table S3: Differences in total exchange (Colo22), related to heatmaps**
For each peptide, the total percent exchange was determined by adding the percent exchange at all timepoints. The difference in total exchange is shown for comparisons that are discussed in the main text. Differences are colored to match how they appear in the related heatmap figures.

| H1 | Activation dynamics[1] | Mutations that *increase* acid stability |
|---|---|---|
| F' Domain$^{N\text{-term}}$ | 17-31 | E31K[2] |
| Vestigial Esterase Domain$^{N\text{-term}}$ | 103-107 | |
| Receptor Binding Domain | 210-218<br>231-236 | N210S[3] |
| Vestigial Esterase Domain$^{C\text{-term}}$ | 267-274 | |
| F' Domain$^{C\text{-term}}$ | 312-324 | |
| Fusion Peptide | 1-8<br>17-21 | |
| F Domain | 55-59<br>140-149<br>151-161 | E47K[4]<br>V55I[5]<br>L99M[6]<br>R106K[7]<br>K153E[5] |
| **H5** | **Activation dynamics (this paper)** | **Mutations that *increase* acid stability** |
| F' domain$^{N\text{-term}}$ | | H18Q[8]<br>K45D[8]<br>K50M[8] |
| Vestigial Esterase Domain$^{N\text{-term}}$ | 62-118 | D104N/I115T[9]<br>H110Y[10] |
| Receptor Binding Domain | 210-219<br>257-268 | K262I[11] |
| Vestigial Esterase Domain$^{C\text{-term}}$ | | G275H[8] |
| F' domain$^{C\text{-term}}$ | 317-328 | H295Q[8]<br>T318I[12] |
| Fusion Peptide | 1-21 | |
| F Domain | 64-80 | K58I[8]<br>N81E[8]<br>E105K[8] |
| **H3** | **Activation dynamics[1]** | **Mutations that *increase* acid stability** |
| F' Domain$^{N\text{-term}}$ | 1-13<br>33-63 | H17Y[13] |
| Vestigial Esterase Domain$^{N\text{-term}}$ | | |
| Receptor Binding Domain | 259-268 | |
| Vestigial Esterase Domain$^{C\text{-term}}$ | | |
| F' Domain$^{C\text{-term}}$ | 315-328 | |
| Fusion Peptide | 1-9 | |
| F Domain | 39-69<br>119-138 | K51(A/E)[13]<br>K58I[14]<br>T156N[15] |

**Table S4: Activation dynamics and acid-stabilizing mutations, related to Figure 8**

Peptides that become more dynamic at low pH in HDX-MS experiments and reported acid-stabilizing mutations.

| Dataset | Viet2004 WT pH 7.4 | Viet2004 WT pH 6.1 | Viet2004 Mutant pH 7.4 | Viet2004 Mutant pH 6.1 | Viet2004 Mutant pH 5.9 |
|---|---|---|---|---|---|
| **HDX reaction details** | 85% D2O buffer, pH* 7.385, labeled at RT. Quenched at pH 2.455 in 2M Urea, 200mM TCEP, 0.2% FA | 85% D2O buffer, pH* 6.072, labeled at RT. Quenched at pH 2.460 in 2M Urea, 200mM TCEP, 0.2% FA | 85% D2O buffer, pH* 7.385, labeled at RT. Quenched at pH 2.455 in 2M Urea, 200mM TCEP, 0.2% FA | 85% D2O buffer, pH* 6.072, labeled at RT. Quenched at pH 2.460 in 2M Urea, 200mM TCEP, 0.2% FA | 85% D2O buffer, pH* 5.875 labeled at RT. Quenched at pH 2.478 in 2M Urea, 200mM TCEP, 0.2% FA |
| **HDX incubation times** | 3 sec, 30 sec, 3.5 min, 30 min | 1 min, 10 min, 1 hr 11 min 10 hr 5 min | 3 sec, 30 sec, 3.5 min, 30 min | 1 min, 10 min, 1 hr 11 min 10 hr 5 min | 1 min 40 sec, 15 min 50 sec, 1 hr 51 min, 15 hr 51 min |
| **HDX controls** | Undeuterated, zero control, fully deuterated | Undeuterated, zero control | Undeuterated, zero control, fully deuterated | Undeuterated, zero control | Undeuterated, zero control, 1 min 40 sec pulse @ 15 hr 51 min timepoint |
| **Standards** | TM-155 TM-151 TM-143 | TM-155 TM-151 TM-143 | TM-155 TM-151 TM-143 | TM-155 TM-151 TM-143 | TM-155 TM-151 TM-143 |
| **Back-exchange** *based on fully deuterated controls | 25 +/- 11% | 25 +/- 11% | 26 +/- 11% | 25 +/- 11% | 25 +/- 11% |
| **Replicates** (biological or technical) | Two technical replicates | Two technical replicates | Two technical replicates | Two technical replicates | Two technical replicates |
| **Repeatability** | stddev of 0.8% across technical replicates | stddev of 0.6% across technical replicates | stddev of 0.7% across technical replicates | stddev of 0.7% across technical replicates | stddev of 0.6% across technical replicates |
| **Dataset** | **Indo2005 WT pH 7.4** | **Indo2005 WT pH 6.1** | **Indo2005 Mutant pH 7.4** | **Indo2005 Mutant pH 6.1** | **Indo2005 Mutant pH 5.8** |
| **HDX reaction details** | 85% D2O buffer, pH* 7.363, labeled at RT. Quenched at pH 2.481 in 2M Urea, 200mM TCEP, 0.2% FA | 85% D2O buffer, pH* 6.082, labeled at RT. Quenched at pH 2.477 in 2M Urea, 200mM TCEP, 0.2% FA | 85% D2O buffer, pH* 7.363, labeled at RT. Quenched at pH 2.481 in 2M Urea, 200mM TCEP, 0.2% FA | 85% D2O buffer, pH* 6.082, labeled at RT. Quenched at pH 2.477 in 2M Urea, 200mM TCEP, 0.2% FA | 85% D2O buffer, pH* 5.840 labeled at RT. Quenched at pH 2.439 in 2M Urea, 200mM TCEP, 0.2% FA |
| **HDX incubation times** | 3 sec, 30 sec, 3.5 min, 30 min | 1 min, 9 min 30 sec, 1 hr 6 min, 9 hr 30 min | 3 sec, 30 sec, 3.5 min, 30 min | 1 min, 9 min 30 sec, 1 hr 6 min, 9 hr 30 min | 1 min 30 sec, 16 min 30 sec, 1 hr 54 min, 16 hr 30 min |

| | | | | | |
|---|---|---|---|---|---|
| **HDX controls** | Undeuterated, zero control, fully deuterated | Undeuterated, zero control | Undeuterated, zero control, fully deuterated | Undeuterated, zero control | Undeuterated, zero control, 1 min 30 sec pulse @ 16 hr 30 min timepoint |
| **Standards** | TM-155 | TM-155 | TM-155 | TM-155 | TM-155 |
| **Back-exchange** *based on fully deuterated controls | 22 +/- 11% | 22 +/- 11% | 22 +/- 11% | 22 +/- 11% | 22 +/- 11% |
| **Replicates** (biological or technical**)** | Two technical replicates | Two technical replicates | Two technical replicates | Two technical replicates | Two technical replicates |
| **Repeatability** | stddev of 0.8% across technical replicates | stddev of 0.7% across technical replicates | stddev of 0.6% across technical replicates | stddev of 0.6% across technical replicates | stddev of 0.8% across technical replicates |
| **Dataset** | **Colo2022 pH 7.4** | **Colo2022 pH 5.8** | | | |
| **HDX reaction details** | 85% D2O buffer, pH* 7.385, labeled at RT. Quenched at pH 2.455 in 2M Urea, 200mM TCEP, 0.2% FA | 85% D2O buffer, pH* 5.828, labeled at RT. Quenched at pH 2.403 in 2M Urea, 200mM TCEP, 0.2% FA | | | |
| **HDX incubation times** | 3 sec, 30 sec, 3.5 min, 30 min | 1 min 50 sec, 18 min 2 hr 6 min, 18 hr | | | |
| **HDX controls** | Undeuterated, fully deuterated | Undeuterated, 1 min 50 sec pulse @ 18 hr timepoint | | | |
| **Standards** | TM-155 TM-151 | TM-155 TM-151 | | | |
| **Back-exchange** *based on fully deuterated controls | 31 +/- 11% | 31 +/- 11% | | | |
| **Replicates** (biological or technical**)** | Two technical replicates | Two technical replicates | | | |
| **Repeatability** | stddev of 0.8% across technical replicates | stddev of 1.0% across technical replicates | | | |

**Table S5: HDX data quality**
HDX reaction details for all HA constructs.
